# Deep learning assisted single particle tracking for automated correlation between diffusion and function

**DOI:** 10.1101/2023.11.16.567393

**Authors:** Jacob Kæstel-Hansen, Marilina de Sautu, Anand Saminathan, Gustavo Scanavachi, Ricardo F. Bango Da Cunha Correia, Annette Juma Nielsen, Sara Vogt Bleshøy, Wouter Boomsma, Tom Kirchhausen, Nikos S. Hatzakis

## Abstract

Sub-cellular diffusion in living systems reflects cellular processes and interactions. Recent advances in optical microscopy allow the tracking of this nanoscale diffusion of individual objects with an unprecedented level of precision. However, the agnostic and automated extraction of functional information from the diffusion of molecules and organelles within the sub-cellular environment, is labor-intensive and poses a significant challenge. Here we introduce DeepSPT, a deep learning framework to interpret the diffusional 2D or 3D temporal behavior of objects in a rapid and efficient manner, agnostically. Demonstrating its versatility, we have applied DeepSPT to automated mapping of the early events of viral infections, identifying distinct types of endosomal organelles, and clathrin-coated pits and vesicles with up to 95% accuracy and within seconds instead of weeks. The fact that DeepSPT effectively extracts biological information from diffusion alone indicates that besides structure, motion encodes function at the molecular and subcellular level.

## INTRODUCTION

Direct observation of cellular processes is now routinely achieved by fluorescence microscopy and single-particle tracking (SPT) techniques^1–8^. These techniques offer the necessary spatiotemporal resolution to localize and track diffusion of individual biomolecules—from small proteins and viruses to organelles or entire cells—in both 2D and 3D environments^4,5,9–12^. The observed diffusion is highly complex and exhibits considerable spatiotemporal and interparticle heterogeneity, reflecting various biological factors such as internalization stages, local environment, oligomerization states, and interactions with elements like the cytoskeleton, membranes, molecular motors, organelles and more^4,10,13,14^. This intrinsic heterogeneity presents a significant analytical challenge and remains a major bottleneck for extracting quantitative insights from single-particle studies (SPT).

Traditionally, extracting diffusional behavior from SPT experiments relies on fitting the mean squared displacement (MSD) ^14–17^, or more recently using machine learning^18–21^. These approaches often transform entire trajectories into a single descriptor like the diffusion coefficient, thus averaging out vital temporal information that is essential to interpret biological processes. Achieving temporal segmentation—a prerequisite for unlocking the rich temporal data inherent in biological processes—is a considerable challenge. For instance, manual annotation requires significant expertise and is prohibitively time-consuming for large datasets, and to date is done primarily in 2D. Methods like Rolling MSD^14–17,22^ and divide-and-conquer^23^ offer automated temporal segmentation, but they are reliant on windowing tracks which introduces a trade-off in temporal sensitivity and accuracy, and they also depend on user-defined, system-specific parameters. Hidden Markov Models^24–27^ can segment traces but only if a diffusional metric, often step length, varies significantly between states^27,28^.

Current state-of-the-art tools based on machine learning^28–34^ can distinguish between diffusional states in 2D with distinct a priori, user-defined diffusional characteristics. While these tools operate well for their specific set of states, they remain less explored for systems displaying multiple, broadly distributed diffusional characteristics common in complex cellular environments and often do not extend to 3D. Achieving temporal segmentation of such heterogeneous behavior is crucial for overcoming the current analytical bottleneck in SPT, yet alone is not enough to decode correlations between heterogeneous behavior, biological context, function, and biomolecular identity.

Identifying biomolecular identity, colocalizing partners, cellular localization, or the time point of a biological event is a challenge in fluorescence microscopy, necessitating specialized analysis, parallelized multicolor and often super-resolution imaging^35–38^. Such experimental design, specialized analysis and fluorescent tagging of biomolecular entities is substantially labor- and material intensive and risks impairing biological function by tagging^39^. These challenges are further compounded by the limitation of using 2-3 imaging channels for quantitative imaging due to spectral overlap. Temporal analysis of diffusional behavior alone could overcome these challenges by acting as an orthogonal probe for extracting biological function, colocalization or identity, minimizing the need for fluorescent tagging, thereby simplifying experimental workflows. However, to date dissecting heterogeneous diffusional behavior remains largely untapped

Here we introduce DeepSPT, a versatile deep learning-based toolbox designed for the rapid, accurate, and automatic temporal analysis of behavior in single-particle tracking (SPT). DeepSPT facilitates extraction of biological insights from 2D or 3D traces solely based on the diffusional characteristics of the tracked objects. The pre-trained DeepSPT pipeline is available as open-source code on GitHub.

## RESULTS

### DeepSPT

DeepSPT is a deep learning framework, encompassing three sequentially connected modules: A temporal behavior segmentation module; a diffusional fingerprinting module; and a task-specific downstream classifier module (Fig. 1a). The first two modules are universal, applicable directly to any trajectory dataset characterized by x, y, (z) and t coordinates across diverse biological systems. The final module capitalizes on experimental data to learn a task that is specific to the system under investigation.

**Fig. 1.**
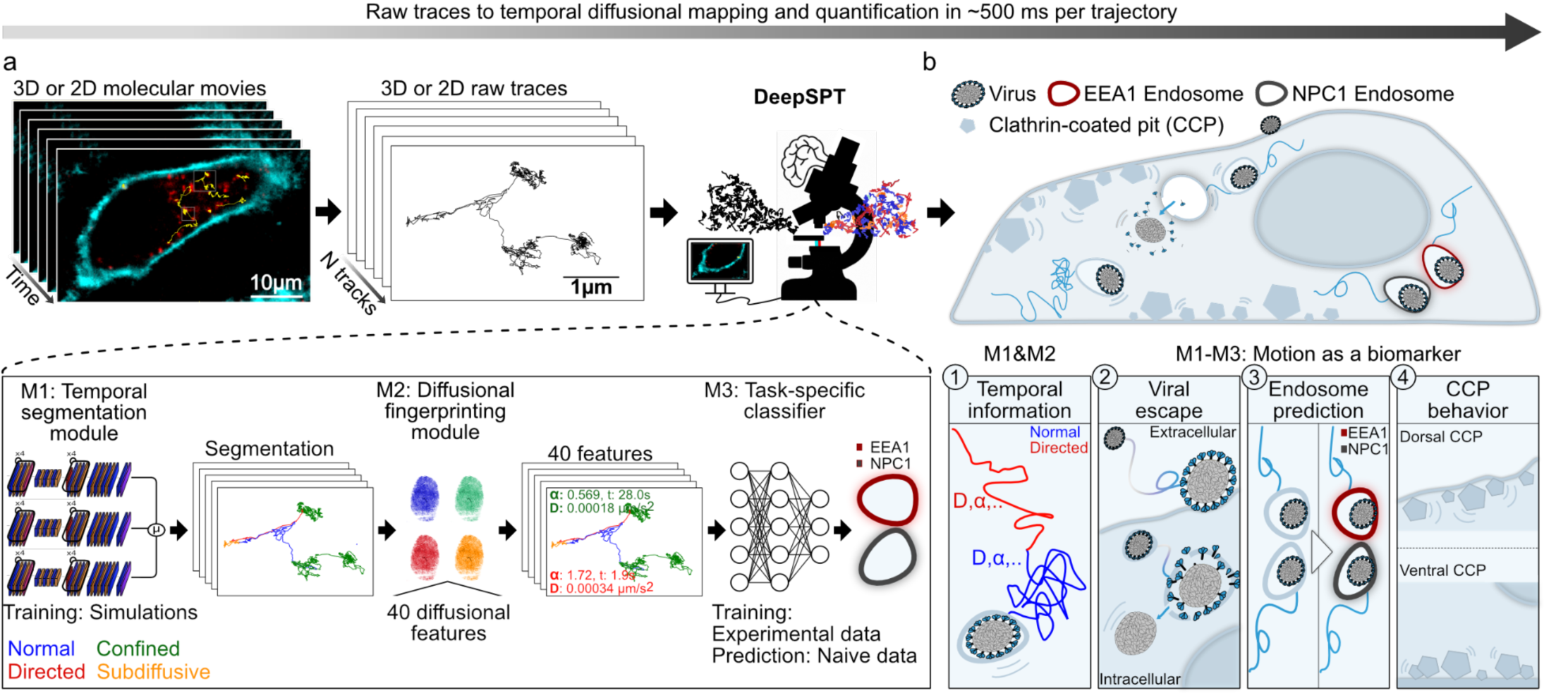
DeepSPT, an agnostic, automated approach for extraction of time-dependent behavior in dynamic systems. **a**, Schematic representation of the DeepSPT pipeline: 2D or 3D molecular movies from fluorescence microscopy imaging produce a set of x, y, (z), t, localizations for each particle yielding a dataset of single particle trajectories. These trajectories are directly fed to DeepSPT consisting of a temporal behavior segmentation module (M1), diffusional fingerprinting module (M2) and a task-specific classifier (M3). The modules of DeepSPT appear in the zoom-in. Firstly, the temporal segmentation module classifies per time point the diffusional behavior (normal, directed, confined or subdiffusive). Secondly, tracks segmented into diffusional behaviors are quantified by multiple diffusional descriptors by the diffusional fingerprinting module. Thirdly, a task-specific classifier trained utilizing the temporal information and the diffusional fingerprints for each track to learn a problem of interest, e.g., identification of endosomal identity based on diffusional behavior of cargo. The entire DeepSPT pipeline has a computational time of ∼500ms per trajectory. **b**, Schematic illustration of selected biological applications enabled by DeepSPT pipeline: 1) Temporal diffusional behavior segmentation, analysis and quantification. 2-4) Applications of DeepSPT to uncover biological insights, based exclusively on diffusional behavior variation. 2) Time point identification of biological events such as detection of viral escape into the cytosol. 3) Prediction of endosomal identity directly using endosomal motion or solely from movement of their cargo. 4) Predicting cellular localization of clathrin-coated pits.

The temporal behavior segmentation module transforms single particle trajectories into sub-segments characterized by distinct diffusional behaviors, processing input directly from x, y, (z), and t coordinates using an ensemble of fully convolutional networks (see Methods). Alongside the predicted diffusional behavior, each time point in the trajectory is assigned a probability estimate for each type of diffusion identified. This study focuses on the four types of diffusional behaviors predominantly reported in biological system^16,25,32,40–42^: 1) Normal diffusion, typifying unhindered random motion; 2) directed motion, as commonly exhibited by molecular motors; 3) confined motion, characterizing limited spaces with reflective boundaries, such as small membranes structures; and 4) subdiffusive motion, indicative of more restrained movement, commonly observed in densely populated cytosolic environments. The module’s training utilized an extensive dataset comprising 900,000 trajectories, exhibiting broadly distributed diffusional properties; this encompasses variations spanning four orders of magnitude in diffusional coefficients, diverse trace durations, varying localization errors, and trajectories displaying multiple, random length diffusional behaviors throughout their lifespan (see Methods). This extensive training set expands DeepSPT’s adaptability across different biological systems and experimental conditions. It is important to note that DeepSPT can be trained to recognize other diffusional attributes and diverse motion types, or to simply predict a uniform global diffusional state in cases of homogenous motion.

The diffusional fingerprinting module within DeepSPT, transforms each identified segment of diffusional behavior into a comprehensive set of 40 descriptive diffusional features, not just encompassing but also expanding upon those enunciated by Pinholt et al^21^. This module serves dual purposes: it facilitates statistical quantification of individual behavior segments for user interpretation; and it generates feature representations crucial for downstream classification tasks (see Methods).

The task-specific downstream classification module of DeepSPT trains and predicts directly on experimental data, which has been transformed to a combined feature set by the temporal and diffusional fingerprinting modules. This module outputs class probability estimates solely utilizing diffusional characteristics for any domain. This is exemplified by predicting important time points during the initial phases of rotavirus infection, differentiating early and late endosomes, and by locating clathrin-coated pits and vesicles to the dorsal or ventral membranes of a cell (Fig. 1b).

### Rapid, Automated Analysis of Temporal Diffusional Behavior

To demonstrate the effectiveness and generalizability of the temporal segmentation capabilities of DeepSPT, we employed three distinct evaluation schemes. First, we used a holdout scheme to assess performance on trajectories withheld during training (Fig. 2a-c). Second, we tested the model’s generalizability using simulated trajectories with a wider distribution in the values of diffusion parameters than those used during training. Third, we compared DeepSPT’s performance against existing state-of-the-art temporal segmentation algorithms.

**Fig. 2.**
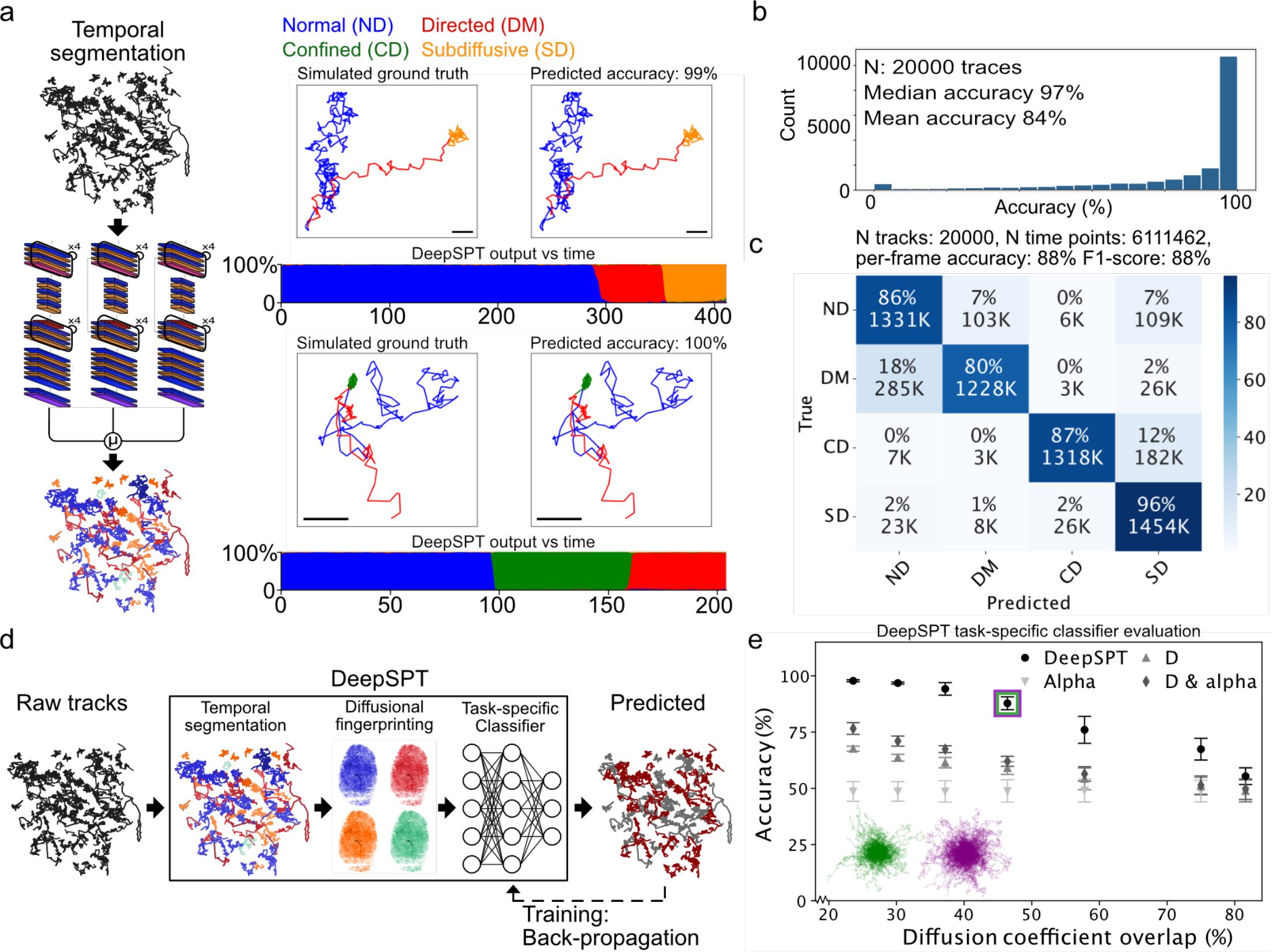
Evaluation of DeepSPT’s temporal behavior segmentation. **a**, Illustration of the temporal segmentation module of DeepSPT along with two examples of DeepSPT prediction on simulated trajectories with heterogeneous diffusion. For representative 3D examples of predictions: Left: 2D projection simulated ground truth color coded to underlying diffusional behavior. Right: Trajectory color coded to DeepSPT’s predictions. Scale bar 500 nm. Bottom: Uncertainty calibrated probability estimates (DeepSPT output vs. time) for each modeled diffusion type per time point, providing transparency into model certainty associated with a given prediction (see Supplementary fig. 3 for uncertainty calibration by temperature scaling). **b**, Histogram of the accuracies associated with each individual 3D trajectory in the test set (N traces: 20.000, N time points: 6111462) and population descriptive statistics such as a median accuracy of 96% (see methods for test set specification). **c**, Confusion matrix based on all predictions (N time points: 6111462) within the 20,000 3D test set trajectories in **b** totaling >6M individual time point predictions. Diagonal entries are correct predictions and off-diagonal indicates confused classes. Each entry reports the absolute number of predictions (K=1000) and its normalization to the actual number of labels in the given class. F1 score of 88% shows DeepSPT to accurately segment and classify heterogeneous diffusion. **d**, Illustration of the DeepSPT classification pipeline. Each raw track is temporally segmented to each of the four diffusional behavior by the segmentation module, transformed into descriptive features by the diffusional fingerprinting module which combines to a unique feature set of temporal and diffusional features which subsequently is fed to a task-specific downstream classifier. **e**, Benchmarking of the DeepSPT classification pipeline against a classifier using MSD features: Diffusion coefficient (D), the anomalous diffusion exponent term (alpha), or both (D & alpha) on simulated data of two classes of trajectories with overlapping diffusional properties (see Methods). Classification accuracy is evaluated at incrementing degrees of overlap in the instantaneous diffusion coefficients. Purple and green trajectories depict trajectories at ∼45% overlap in diffusion coefficients indicated by the purple and green boxes. DeepSPT significantly outperforms all three MSD features up to 75% overlap in diffusion coefficient (all p-values < 0.001 using two-sided Welch t-test, N=5 per condition) and at 82% overlap DeepSPT significantly outperforms D and alpha (all p-values < 0.05).

In the holdout validation, we assessed the temporal segmentation on a test set comprising 20,000 simulated trajectories, of which 80% exhibited heterogeneous motion and 20% showed homogeneous motion (see Methods). The trajectories spanned a broad range of diffusional parameters and four motion types (Fig.2a, see Methods, Supplementary fig. 4). DeepSPT not only accurately identified temporal changepoints (Fig. 2a) but also yielded time-resolved probability estimates that may serve as an adjustable post-processing parameter (Fig. 2a, DeepSPT output vs. time). We calibrated these probability estimates using temperature scaling^43^ (Supplementary fig. 3) to enhance reliability and mitigate overconfidence, Quantification of DeepSPT’s classification performance revealed a median accuracy of 96% per trace and 84% mean accuracy per frame for all four motion types (Fig. 2b,c, and Supplementary fig. 1). The model achieved 91% mean accuracy for three motion types—normal, directed, and confined/subdiffusive—and 97% for two motion types normal/directed versus confined/subdiffusive motion types (Supplementary fig. 1); and 91% for homogeneous motion (Supplementary fig. 2). DeepSPT achieved an F1 value of 88% for both 3D and 2D data sets (Fig. 2c, Supplementary fig. 1). In all cases it has an inference time of less than 40 ms per trajectory. Subdiffusive motion was classified with 96% accuracy, directed motion with 80%, normal and confined motion with 86% and 87% accuracy, respectively. Minimal confusion existed between dissimilar motion types, highlighting DeepSPT’s capability to differentiate between restricted and free motion types. This strength became more apparent when the model was tasked with identifying fewer motion categories (Supplementary fig. 1). Robust performance of DeepSPT was further confirmed across a variety of diffusional properties, track durations, and localization errors, even for parameter ranges not included in the training set. DeepSPT excelled for traces longer than 20 frames and localization errors equal or smaller than the actual diffusional step lengths (Supplementary fig. 8-10)., demonstrating its adaptability to various experimental setups.

We benchmarked DeepSPT’s ability to segment heterogeneous diffusion against a high-performing LSTM-based method^31^ and the widely used rolling MSD approach^14–17,22^ (Fig. 1c). We focused on these methods over others, such as HMM^24–27^, because they do not require large variations in step lengths to detect changes in diffusional behavior (see Supplementary fig. 11,12). Each method was tested on 2D trajectories, given the LSTM-based technique’s 2D limitation. The LSTM-based method achieved classification accuracies of 44%, 58%, and 72%, outperforming the rolling MSD’s 34%, 51%, and 65% for four, three, and two diffusional behaviors, respectively. DeepSPT, in contrast, attained accuracies of 88%, 91%, and 97% for the same categories, outperforming current state-of-the-art and highlighting its improved competence in agnostic segmentation and classification of heterogeneous diffusion.

We then qualitatively evaluated DeepSPT on 2D experimental data sets (Supplementary fig. 6,7). Specifically, for human insulin we labeled with Atto-655 and recorded its spatiotemporal localization in HeLa cells using 2D using live-cell spinning disk confocal fluorescence microscopy (see Methods). Using DeepSPT we report insulin intracellular transport mainly exhibited subdiffusive behavior but included segments of directed motion. The directed motion aligns with motor-protein diffusion patterns indicative of active cellular trafficking, establishing DeepSPT as a potential tool for studying transport mechanisms across diverse experimental contexts.

Classification of biomolecular identity requires an addition to temporal segmentation. We combine all modules of DeepSPT to demonstrate its capabilities to leverage subtle diffusional variations to classify heterogeneous behavior. The integration of DeepSPT’s segmentation and fingerprinting modules allows for the transformation of any trajectory into a feature representation containing both temporal and diffusional features, which can then be fed to a downstream classifier (Fig. 2d). To demonstrate the descriptive power of DeepSPT’s integrated approach, we assessed the classification performance of DeepSPT on two classes of simulated trajectories (1000 tracks) with overlapping diffusional properties (see Methods). Keeping all diffusional features except the diffusion coefficient constant, we evaluated the classification accuracy by stratified 5-fold cross-validation for varying degrees of overlap in diffusion coefficients between the classes (see Methods). DeepSPT achieved up to 98% accuracy and maintained 76% accuracy even when the overlap in diffusion coefficients was around 57%. This performance significantly (by Welch t-test, fig. 2e) outperformed that of basic MSD features, which attained accuracies ranging from approximately 49% to 76% (Fig. 2e). Such robust classification highlights DeepSPT’s ability to discern and exploit subtle differences in diffusional properties.

### DeepSPT accelerates detection of viral escape using motion as a marker

We validated the operational efficacy of DeepSPT to extract information from 3D live-cell SPT data of rotavirus (Fig. 3a, see Methods). The entry process of rotavirus into cells, the first step for infection, involves glycolipid-mediated membrane association of the virus, vesicular engulfment and internalization, virus-initiated membrane permeabilization, calcium-dependent uncoating of outer proteins, membrane disruption, and cytosolic delivery of the viral genome for subsequent RNA production (Fig. 3a)^44^. Previous single-particle tracking in BSC-1 cells using confocal imaging indicated that the uncoating step correlates with a change in diffusional behavior, hinting motion as a potential marker of biological behavior^44,45^.

**Fig. 3.**
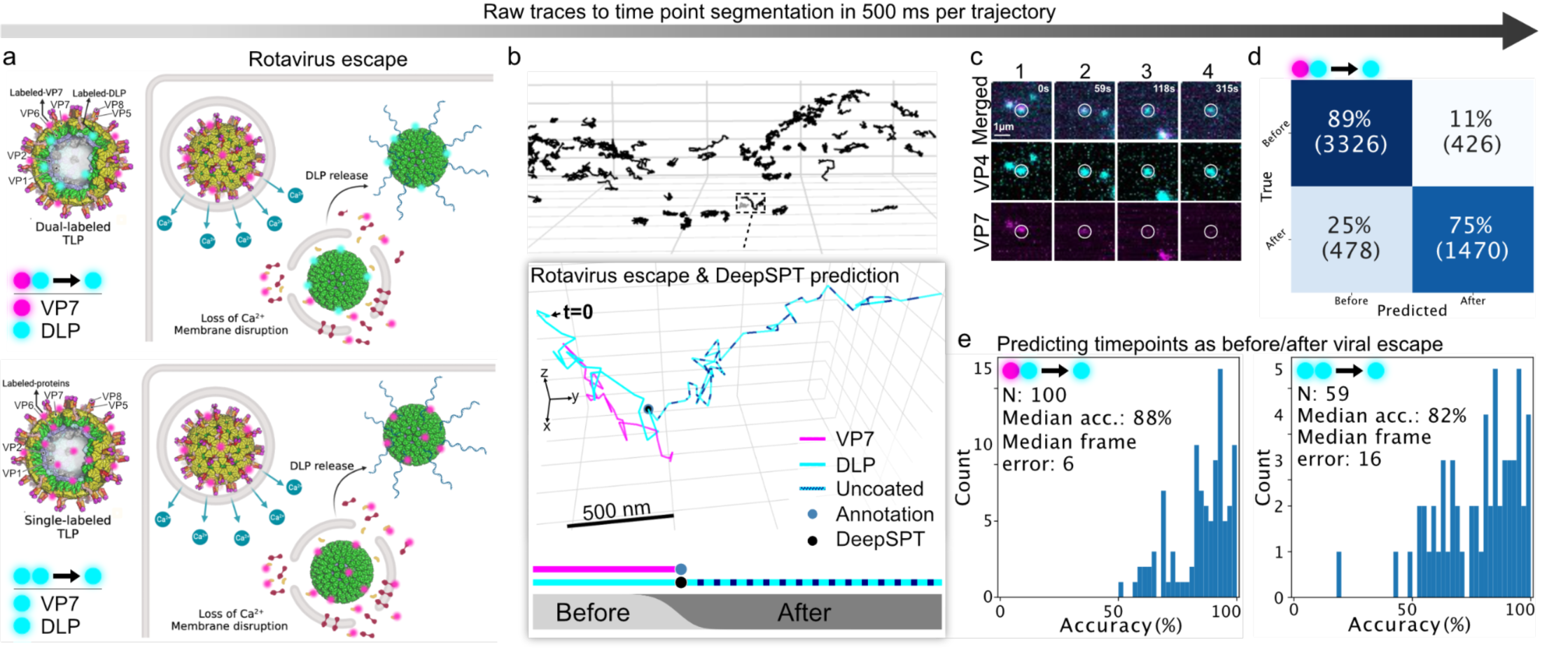
Rapid and precise classification of rotavirus uncoating by DeepSPT based exclusively on diffusional behavior. **a**, Schematic illustration of typical stages of rotavirus cell-entry pathway from interactions at the plasma membrane, membrane engulfment, membrane permeabilization, calcium-dependent uncoating and escape to the cytosol where RNA production can begin. Top and bottom panel display the two experimental approaches used for single rotavirus tracking: Top, Dual-labeled by recombinant construction of rotavirus with fluorescently tagged DLP and VP7. Bottom, monochromatic, chemical labeling of free lysines by Atto560. **b**, 3D tracks of individual rotavirus particles in a single cell acquired by live-cell lattice light sheet microscopy (LLSM). Zoom-in: Example of rotavirus by parallel multicolor imaging of DLP and VP7 (see Methods). The time point for loss of VP7 signal, indicative of uncoating and viral escape (blue dot), is correctly identified by DeepSPT (black dot). Bottom Insets: 1D representation of DLP and VP7 signal with annotations for loss of VP7 and DeepSPT prediction. Softmax output, providing time-resolved probability estimates of “before uncoating” and “after uncoating”. **c**, Sum intensity projections of the 3D live-cell LLSM raw data from a region of interest surrounding the track in the zoom-in. The insets contain parallel imaging of DLP (cyan) and VP7 (magenta). Numbered columns showing different observed stages of the virus’s lifetime for DLP and VP7 from colocalization to uncoating. **d**, Confusion matrix displaying DeepSPT classification performance of predicting time points as “before uncoating” or “after uncoating” as compared to ground truth colocalization analysis, entries normalized to true labels for dual-labeled rotavirus (see Methods) (top left: True before, top right: False after, bottom left: False before, bottom right: True after). **e-f**, Histogram of DeepSPT classification accuracies as percentage of time points correctly predicted “before uncoating” or “after uncoating” in individual tracks. **e**, Dual-labeled rotavirus showing median accuracy 88% (100 tracks, N=1 coverslip experiments, 4 movies). **f**, monochromatically labeled rotavirus showing median accuracy 82% (59 tracks, N=5 coverslip experiments, 13 movies). DeepSPT requires 500 ms processing time per trajectory to transit from raw trajectories to unique feature representations to classification of time points (classification time is ∼1 millisecond), thus, accelerating the analysis by a minimum of 4 orders of magnitude as compared to manual annotations.

To test DeepSPT’s capacity to detect the uncoating and cytosolic delivery events using only motion, we used 3D live-cell lattice light sheet microscopy^5,46^ (LLSM) to image cell entry of reconstituted rotavirus^44,47^ labeled with either: Atto560 on VP7, an outer shell protein, and Atto642 on DLP, the double layered particle; or with Atto560 on the entire virus including both VP7 and DLP (see Fig. 3a,b). Trajectories captured via LLSM (see Fig. 3b) underwent temporal segmentation and diffusional fingerprinting using DeepSPT’s modules in rolling windows for sequential representation (see Methods). These processed trajectories were then classified as either “before uncoat” or “after uncoat” through a sequence-to-sequence based model, transforming coordinates in time into time-resolved predictions, serving as the task-specific classifier (see Methods). The ground truth for uncoating events for dual-labeled rotavirus was established by determining the extent of colocalization of differentially labeled DLP and VP7 (see Methods). A representative example of rotavirus uncoating alongside DeepSPT’s consistent prediction and snapshots of the raw data are shown (Fig. 3b zoom-in and Fig. 3c).

DeepSPT correctly identified 89% of “pre-uncoating” and 75% of “post-uncoating” time points, yielding a mean accuracy of 85% and a median accuracy of 88% across 100 dual-labeled rotavirus trajectories. This high level of accuracy translated to a median error of just 6 frames in determining the uncoating time point (Fig. 3d,e). Unlike traditional methods, which often require manual analysis taking several minutes to hours per trajectory, DeepSPT automated the identification process, reducing the time to milliseconds per viral trajectory offering massive acceleration and minimizing any human bias.

Importantly, DeepSPT outputs these predictions, in 500 ms per trajectory, based solely on the motion captured in the DLP trajectories, rendering the secondary VP7 channel redundant. When tested on rotavirus labeled with the same fluorophore on both DLP and VP7 (see Methods), DeepSPT exhibited similar performance, achieving a median accuracy of 82% and a mean accuracy of 78% (see Methods, Fig. 3f). It’s worth noting that it required labor-intensive (∼8 working days) manual annotation to acquire the ground truth annotations based on intensity loss of the 560 nm channel (see Supplementary fig. 13) while taking less than a minute for DeepSPT. By using motion as a marker for viral uncoating, DeepSPT simplifies experimental design and preparation, avoiding the need for constructing dual-labeled viruses. Thus DeepSPT frees up one of the 2-3 available imaging channels, thereby increasing the information content in fluorescence microscopy experiments. To the best of our knowledge, these results constitute the first instance of detecting viral escape into the cytosol based solely on motion, and without the need for multicolor labeling.

### DeepSPT enables label-minimal colocalization analysis and identifies cellular localization through motion

The capacity to identify biomolecular identity, colocalization partners or to infer subcellular localization based solely on diffusional properties could minimize the need for multicolor imaging and the labor-intensive efforts associated with the creation of cell lines expressing the relevant fluorescent cellular markers. Early and late endosomes for example are difficult to differentiate without multicolour labeling as they might appear to exhibit similar dimensions, are distributed with similar spatial density, and display nearly identical diffusion coefficients^49–51^. Traditionally, their identification requires labeling of each endosomal type through antibodies specific for endogenous protein markers enriched in a given type of endosome, or by ectopic expression of these markers. Based on multicolour labeling, efforts have been made using population-based multiparametric image analysis of internalized cargo distribution and compartment morphology in fixed samples to deduce general principles of the endocytic machinery^48^.

Here we assess whether DeepSPT can determine endosomal identity based solely on diffusional characteristics, reducing the need for multicolour labeling (Fig. 4a). We used two-color live-cell LLSM to track early endosomes endogenously tagged by gene editing with (EEA1-mScarlett) and late endosomes tagged with (NPC1-Halo-JFX646). Their trajectories display indistinguishable diffusion coefficients and alpha values (Fig. 4b), as well as similar diffusional behavior propensities (Supplementary fig. 14) challenging endosomal identity prediction. To address this issue, we introduced a decision confidence threshold, requiring the classifier’s probability estimate to surpass this threshold for prediction acceptance^35^. In a 10-fold stratified cross-validation scheme with varying decision confidence thresholds, DeepSPT achieved accuracies ranging from 70±1.3% to 82±1.8% in classifying EEA1-positive from NPC1-positive compartments. Increasing the confidence threshold improved accuracy but reduced the number of accepted tracks (Fig. 4c). At a 60% confidence threshold, DeepSPT identified endosomal types with an accuracy of 72±1.4% (Fig. 4d), and a recall of 72±3% for EEA1-positive compartments and 72±1.4% for NPC1-positive compartments (Fig. 4d). DeepSPT significantly outperformed the commonly used MSD analysis that reached accuracies of 48±4%, 55±1.6 and 60±1.4% (Supplementary fig. 15,16) in endosomal classification by using the variation in alpha values, variation in the diffusion coefficient (D), or combining both alpha and D, respectfully. DeepSPT, solely using the diffusion traits of endosomal cargo, achieved 91% and 94% of the recall observed in direct prediction of EEA1-positive and NPC1-positive compartments, respectively (Fig. 4e). The ability of DeepSPT to differentiate early from late endosomes based solely on their motion or that of their cargo, accelerates data acquisition and analysis while minimizing potential perturbations or/and the need for multicolour tagging.

**Fig. 4.**
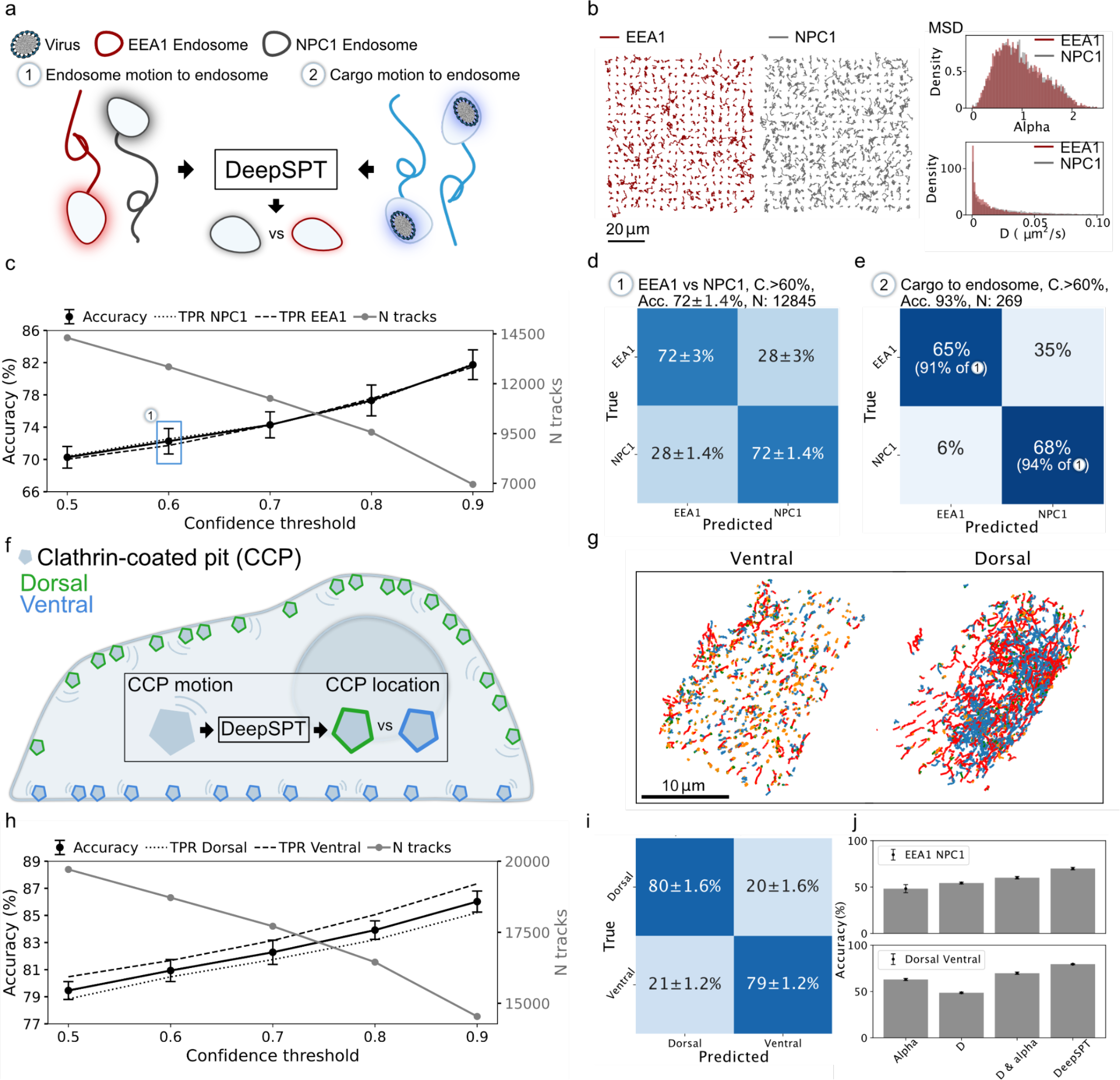
Prediction of endosomal identity and AP2 cellular localization based exclusively on temporal variations in diffusional behavior. **a**, Illustration of endosomal identity prediction solely on the diffusional properties of the endosomal marker or by the motion of endosomal cargo. **b**, Left 2D projection of randomly sampled trajectories of EEA1-mScarlett-(dark red) and NPC1-Halo-JFX646-(gray) positive endosomes acquired by 3D LLSM revealing visually similar trajectories. Right: Distribution of alpha anomalous coefficient (top) and diffusion coefficient (bottom) for EEA1-positive (dark red) and NPC1-positive (gray) displaying practically indiscernible distributions. **c**, Twin axes plot of accuracy and true positive rate (TPR) (left) as well as number of tracks (right) versus confidence threshold (see Methods) of the full classification pipeline of DeepSPT (Fig. 2d). Increasing the confidence threshold enhances accuracy and reduces the number of accepted traces. **d**, Confusion matrices of DeepSPT classification accuracy on prediction of EEA1-positive vs NPC1-positive compartments showing accuracy of 72+-1.4% with a confidence threshold at 60% (top left: True EEA1, top right: False NPC1, bottom left: False EEA1, bottom right: True NPC1). Data is from (N=4 coverslip experiments, 35 movies). **e**, Confusion matrix of DeepSPT classification accuracy on prediction of EEA1-positive vs NPC1-positive compartments now using trajectories of their cargo, i.e., rotavirus with the confidence threshold at 60%. Results in parenthesis show accuracies normalized to results of (d) to compare to the direct prediction accuracy of endosomes. Data consists of 269 tracks (N=12 coverslip experiments, 44 movies). **f**, Illustration of prediction of AP2 complexes’ cellular localization by DeepSPT based exclusively on their diffusional behavior. **g**, 2D projections of AP2 trajectories below and above 500 nm from the coverslip. All trajectories spending >20% of their lifetime above the 500 nm are considered dorsal, while the rest are considered ventral. Trajectories are color-coded by DeepSPT segmentation of diffusional behavior. **h**, Twin axes plot of accuracy (left) and true positive rate (TPR) as well as number of tracks (right) versus confidence threshold (see Methods) of the full classification pipeline of DeepSPT for AP2 data. **i**, Confusion matrices of DeepSPT classification accuracy on prediction of AP2 showing accuracy of 79.5+-0.6% at 50% confidence threshold. Data consists of 19712 tracks from (N=5 coverslip experiments, 13 movies). **j**, Benchmark of using DeepSPT versus conventional metrics based on MSD features: Diffusion coefficient (D) or the anomalous diffusion exponent term (alpha) for both EEA1-positive compartments vs NPC1-positive (EEA1NPC1) and dorsal AP2 vs ventral AP2. DeepSPT significantly outperforms common MSD analysis (all p-values < 0.00001 using two-sided Welch t-test, N=10 per condition).

To assess the universality of DeepSPT to infer identity, we applied it to a new dataset of single-particle trajectories of the assembly of clathrin-coated pits and coated vesicles forming at the cell surface. The dynamic assembly and intracellular location of these structures was obtained by tracking the clathrin AP2 adaptor complex, gene-edited at its sigma subunit with eGFP, using 3D live LLSM. A 2D projection of the acquired AP2 trajectories qualitatively indicated that the diffusional properties of AP2 were correlated with cellular location^52^ —dorsal versus ventral cell surface (Fig. 4f,g). This correlation was quantitatively confirmed by DeepSPT’s temporal segmentation of the 3D traces (Supplementary Fig. 14). DeepSPT accurately predicted the cellular location of AP2 in a 10-fold cross-validation scheme, yielding accuracies that varied from 79.5±0.6% to 86.0±0.8% at different confidence thresholds (Fig. 4h). Without applying any confidence filter (i.e., a 50% threshold), DeepSPT classified the cellular location of AP2 with recalls of approximately 80% for both classes (Fig. 4i). In contrast, pinpointing the cellular location of AP2 using MSD features reached accuracies of 62.5±1.8% 48.7±0.8% and 70±1.3%, with a maximum recall for dorsal tracks of 60±2% (Fig. 4g, Supplementary fig. 15,16). Subtle diffusional variations across systems, while missed by common tools, are utilized by DeepSPT to precisely output biological information in complex systems. DeepSPT achieves this across various biological contexts, imaging protocols, and experimental conditions.

## Discussion

The diffusion of biomolecules within cells exhibits both spatial and temporal heterogeneity and varies across biological systems and functionalities but extracting quantitative temporal information from live-cell imaging is currently an analytical bottleneck and often relies on system-specific analysis or even manual annotation. DeepSPT overcomes this bottleneck by providing a universal framework to transition from raw trajectories to quantitative temporal information rapidly precisely and with minimal human intervention both 2D and 3D diffusion. Trained on trajectories with broadly distributed diffusional properties, DeepSPT consistently outperformed existing state-of-the-art toolboxes both in segmenting and classifying diverse heterogeneous diffusional behaviors, both in simulated and experimental data. The implementation of uncertainty-calibrated probability estimates enhances the transparency of DeepSPT’s output, enabling users with limited a priori knowledge of the biological system to gauge model certainty. The minimal requirement for human intervention highlights DeepSPT’s potential to enhance both the reproducibility and robustness of conclusions across different laboratories. Being open-source and freely available to the public, allows future users to perform customized analyses according to individual research needs.

The precise temporal segmentation combined with the comprehensive quantification of diffusional properties of DeepSPT, coupled with its trained downstream classifier, facilitate rapid prediction of viral uncoating events—achieving results in seconds as opposed to weeks as required for manual annotation. This four orders of magnitude acceleration, not only marks the first deep-learning-assisted identification of viral uncoating but also shifts the bottleneck in single-particle discoveries from data analysis to data acquisition. It even introduces the potential for virtually real-time analysis of early stages of viral infection.

Subtle diffusional variations in 2D or 3D, while missed by common tools, are utilized by DeepSPT to precisely output biological information in complex systems across various biological contexts, imaging protocols, and experimental conditions. For example, DeepSPT discerned EEA1-positive from NPC1-positive compartments solely based on their respective 3D diffusional characteristics, or that of their cargo with accuracies of 72% significantly outperforming the commonly used MSD analysis that reached accuracies of 50-60%. These findings prompt further mechanistic studies to explore whether divergent diffusional behaviors stem from distinct external interaction partners, inherent physical differences between endosomal compartments, or other variables. DeepSPT similarly pinpointed the cellular location of AP2 on 3D data with an accuracy of 80%, significantly outperforming common analysis reaching ∼50-70% accuracy. The distinct diffusional behaviors of AP2 highlighted the importance of careful selection in imaging setups. Applied on 2D data of insulin internalization DeepSPT found insulin mainly exhibits subdiffusive behavior but included segments of directed motion indicative of active transport. DeepSPT’s ability to accurately quantify heterogeneous behaviors in both 2D and 3D, across diverse biological systems and under varying experimental and imaging conditions, attests to its utility as a universal platform for characterizing heterogeneous diffusion across systems.

DeepSPT’s capacity for predicting viral uncoating events, identifying endosomal types, and discerning colocalization partners and cellular localization solely based on diffusion extends the traditional structure-to-function paradigm in proteins to a novel motion-to-function paradigm. This suggests that, alongside structure^9,53–55^, motion can also serve as an indicator of both function and identity. This development opens avenues for employing motion as a biomarker and for label-minimal analyses—effectively substituting fluorescent labels with temporal diffusional analysis. Such a shift could simplify experimental design and reduce preparation time, or potentially enrich experiments by reallocating redundant fluorescent markers for other applications.

Widespread implementation of DeepSPT across laboratories could facilitate the creation of comprehensive libraries detailing characteristic movements of cells, subcellular structures and biomolecules. An open-source diffusional library of this kind would offer a new instrument for the scientific community, aiding in the exploration of 4D cell biology through temporal diffusional behavior.

## Author contributions

J.K.H., N.S.H. and T.K. wrote the paper with feedback from all the authors. J.K.H. performed all computational work with input from T.K. W.B. and N.S.H. M.D.S. performed all rotavirus preparation. A.S., R.D.C.C. and G.S. performed LLSM imaging with the presence of JKH at T.K. laboratory at Harvard Medical School. A.J.J., and S.B. performed SDCM imaging and insulin assays at N.S.H. laboratory at University of Copenhagen. J.K.H. and N.S.H. conceived the project idea. N.S.H. had the overall project management with tight interactions with T.K.

## Acknowledgement

We thank members of our laboratories for help and encouragement. We thank Narendra Kumar Mishra and Professor Knud Jørgen Jensen for discussion concerning the preparation and use of insulin for our experiments. This work was funded by the Villum Foundation by being part of BioNEC (grant 18333) to J.K.H. and N.S.H., the Novo Nordisk foundation challenge center for Optimised Oligo escape (T.K., co-PI), the center for 4D cellular dynamics N.S.H. is affiliated with The Novo Nordisk Foundation Center for Protein Research (CPR) funded by a generous donation from the Novo Nordisk Foundation (grant no. NNF14CC0001). J.K.H., A.J.J., S.B., and N.S.H. are members of the Integrative Structural Biology Cluster (ISBUC) at the University of Copenhagen. VILLUM FONDEN (40516) for W.B. and N.S.H. The Novo Nordisk Foundation Center for Basic Machine Learning Research in Life Science (NNF20OC0062606) for W.B. T.K. acknowledges support from NIH Maximizing Investigators’ Research Award (MIRA) GM130386, NIH Grant AI163019, 1R01, IONIS Pharmaceuticals. M.D.S was supported by NIH/NCI CA13202 grant to S.C.H. Figure 3a was created with BioRender.com.

## Notes

### Competing Interest Statement

The authors have declared no competing interest.

## Bibliography

1. Cocucci, E., Aguet, F., Boulant, S. & Kirchhausen, T. The first five seconds in the life of a clathrin-coated pit. Cell 150, 495–507 (2012).

2. He, K. et al. Dynamics of phosphoinositide conversion in clathrin-mediated endocytic traffic. Nature 552, 410–414 (2017).

3. Sungkaworn, T. et al. Single-molecule imaging reveals receptor-G protein interactions at cell surface hot spots. Nature 550, 543–547 (2017).

4. Johnson, C., Exell, J., Lin, Y., Aguilar, J. & Welsher, K. D. Capturing the start point of the virus-cell interaction with high-speed 3D single-virus tracking. Nat. Methods 19, 1642–1652 (2022).

5. Liu, T.-L. et al. Observing the cell in its native state: Imaging subcellular dynamics in multicellular organisms. Science 360, (2018).

6. Thomsen, R. P. et al. A large size-selective DNA nanopore with sensing applications. Nat. Commun. 10, 5655 (2019).

7. Aguet, F. et al. Membrane dynamics of dividing cells imaged by lattice light-sheet microscopy. Mol. Biol. Cell 27, 3418–3435 (2016).

8. Moses, M. E. et al. Single-Molecule Study of Thermomyces lanuginosus Lipase in a Detergency Application System Reveals Diffusion Pattern Remodeling by Surfactants and Calcium. ACS Appl. Mater. Interfaces 13, 33704–33712 (2021).

9. Jensen, S. B. et al. Biased cytochrome P450-mediated metabolism via small-molecule ligands binding P450 oxidoreductase. Nat. Commun. 12, 2260 (2021).

10. Gabriele, M. et al. Dynamics of CTCF- and cohesin-mediated chromatin looping revealed by live-cell imaging. Science 376, 496–501 (2022).

11. Chenouard, N. et al. Objective comparison of particle tracking methods. Nat. Methods 11, 281–289 (2014).

12. Jaqaman, K. et al. Robust single-particle tracking in live-cell time-lapse sequences. Nat. Methods 5, 695–702 (2008).

13. Wan, F. et al. Ultrasmall TPGS-PLGA Hybrid Nanoparticles for Site-Specific Delivery of Antibiotics into Pseudomonas aeruginosa Biofilms in Lungs. ACS Appl. Mater. Interfaces 12, 380–389 (2020).

14. Dahan, M. et al. Diffusion dynamics of glycine receptors revealed by single-quantum dot tracking. Science 302, 442–445 (2003).

15. Gal, N., Lechtman-Goldstein, D. & Weihs, D. Particle tracking in living cells: a review of the mean square displacement method and beyond. Rheol. Acta 52, 425–443 (2013).

16. Arcizet, D., Meier, B., Sackmann, E., Rädler, J. O. & Heinrich, D. Temporal analysis of active and passive transport in living cells. Phys. Rev. Lett. 101, 248103 (2008).

17. Michalet, X. Mean square displacement analysis of single-particle trajectories with localization error: Brownian motion in an isotropic medium. Phys. Rev. E Stat. Nonlin. Soft Matter Phys. 82, 041914 (2010).

18. Granik, N. et al. Single-Particle Diffusion Characterization by Deep Learning. Biophys. J. 117, 185–192 (2019).

19. Kinder, M. & Brauer, W. Classification of trajectories—Extracting invariants with a neural network. Neural Netw. 6, 1011–1017 (1993).

20. Kowalek, P., Loch-Olszewska, H. & Szwabiński, J. Classification of diffusion modes in single-particle tracking data: Feature-based versus deep-learning approach. Phys. Rev. E 100, 032410 (2019).

21. Pinholt, H. D., Bohr, S. S.-R., Iversen, J. F., Boomsma, W. & Hatzakis, N. S. Single-particle diffusional fingerprinting: A machine-learning framework for quantitative analysis of heterogeneous diffusion. Proc Natl Acad Sci USA 118, (2021).

22. Benning, N. A. et al. Dimensional Reduction for Single-Molecule Imaging of DNA and Nucleosome Condensation by Polyamines, HP1α and Ki-67. J. Phys. Chem. B 127, 1922–1931 (2023).

23. Vega, A. R., Freeman, S. A., Grinstein, S. & Jaqaman, K. Multistep track segmentation and motion classification for transient mobility analysis. Biophys. J. 114, 1018–1025 (2018).

24. Monnier, N. et al. Inferring transient particle transport dynamics in live cells. Nat. Methods 12, 838–840 (2015).

25. Persson, F., Lindén, M., Unoson, C. & Elf, J. Extracting intracellular diffusive states and transition rates from single-molecule tracking data. Nat. Methods 10, 265–269 (2013).

26. Slator, P. J., Cairo, C. W. & Burroughs, N. J. Detection of Diffusion Heterogeneity in Single Particle Tracking Trajectories Using a Hidden Markov Model with Measurement Noise Propagation. PLoS ONE 10, e0140759 (2015).

27. Chen, Z., Geffroy, L. & Biteen, J. S. NOBIAS: Analyzing anomalous diffusion in single-molecule tracks with nonparametric Bayesian inference. Front. Bioinform. 1, (2021).

28. Arts, M., Smal, I., Paul, M. W., Wyman, C. & Meijering, E. Particle mobility analysis using deep learning and the moment scaling spectrum. Sci. Rep. 9, 17160 (2019).

29. Dosset, P. et al. Automatic detection of diffusion modes within biological membranes using back-propagation neural network. BMC Bioinformatics 17, 197 (2016).

30. Matsuda, Y., Hanasaki, I., Iwao, R., Yamaguchi, H. & Niimi, T. Estimation of diffusive states from single-particle trajectory in heterogeneous medium using machine-learning methods. Phys. Chem. Chem. Phys. 20, 24099–24108 (2018).

31. You, B. & Yang, G. Attention-based LSTM for Motion Switching Detection of Particles in Living Cells. in 2021 International Joint Conference on Neural Networks (IJCNN) 1–6 (IEEE, 2021). doi:10.1109/IJCNN52387.2021.9533629.

32. Wagner, T., Kroll, A., Haramagatti, C. R., Lipinski, H.-G. & Wiemann, M. Classification and segmentation of nanoparticle diffusion trajectories in cellular micro environments. PLoS ONE 12, e0170165 (2017).

33. Helmuth, J. A., Burckhardt, C. J., Koumoutsakos, P., Greber, U. F. & Sbalzarini, I. F. A novel supervised trajectory segmentation algorithm identifies distinct types of human adenovirus motion in host cells. J. Struct. Biol. 159, 347–358 (2007).

34. Muñoz-Gil, G. et al. Objective comparison of methods to decode anomalous diffusion. Nat. Commun. 12, 6253 (2021).

35. Thomsen, J. et al. DeepFRET, a software for rapid and automated single-molecule FRET data classification using deep learning. eLife 9, (2020).

36. Ejdrup, A. L. et al. A density-based enrichment measure for assessing colocalization in single-molecule localization microscopy data. Nat. Commun. 13, 4388 (2022).

37. Dunn, K. W., Kamocka, M. M. & McDonald, J. H. A practical guide to evaluating colocalization in biological microscopy. Am J Physiol, Cell Physiol 300, C723–42 (2011).

38. Malle, M. G. et al. Single-particle combinatorial multiplexed liposome fusion mediated by DNA. Nat. Chem. 14, 558–565 (2022).

39. Merino Urteaga, R. & Ha, T. Mind your tag in single-molecule measurements. Cell Rep. Methods 3, 100623 (2023).

40. Monnier, N. et al. Bayesian approach to MSD-based analysis of particle motion in live cells. Biophys. J. 103, 616–626 (2012).

41. Saxton, M. J. & Jacobson, K. Single-particle tracking: applications to membrane dynamics. Annu. Rev. Biophys. Biomol. Struct. 26, 373–399 (1997).

42. Ruthardt, N., Lamb, D. C. & Bräuchle, C. Single-particle tracking as a quantitative microscopy-based approach to unravel cell entry mechanisms of viruses and pharmaceutical nanoparticles. Mol. Ther. 19, 1199–1211 (2011).

43. Guo, C., Pleiss, G., Sun, Y. & Weinberger, K. Q. On Calibration of Modern Neural Networks. arXiv (2017) doi:10.48550/arxiv.1706.04599.

44. Abdelhakim, A. H. et al. Structural correlates of rotavirus cell entry. PLoS Pathog. 10, e1004355 (2014).

45. Salgado, E. N., Garcia Rodriguez, B., Narayanaswamy, N., Krishnan, Y. & Harrison, S. C. Visualization of Calcium Ion Loss from Rotavirus during Cell Entry. J. Virol. 92, (2018).

46. Chen, B.-C. et al. Lattice light-sheet microscopy: imaging molecules to embryos at high spatiotemporal resolution. Science 346, 1257998 (2014).

47. Aoki, S. T. et al. Cross-linking of rotavirus outer capsid protein VP7 by antibodies or disulfides inhibits viral entry. J. Virol. 85, 10509– 10517 (2011).

48. Collinet, C. et al. Systems survey of endocytosis by multiparametric image analysis. Nature 464, 243–249 (2010).

49. Rink, J., Ghigo, E., Kalaidzidis, Y. & Zerial, M. Rab conversion as a mechanism of progression from early to late endosomes. Cell 122, 735– 749 (2005).

50. Gruenberg, J. & van der Goot, F. G. Mechanisms of pathogen entry through the endosomal compartments. Nat. Rev. Mol. Cell Biol. 7, 495– 504 (2006).

51. Piper, R. C. & Katzmann, D. J. Biogenesis and function of multivesicular bodies. Annu. Rev. Cell Dev. Biol. 23, 519–547 (2007).

52. Cocucci, E., Gaudin, R. & Kirchhausen, T. Dynamin recruitment and membrane scission at the neck of a clathrin-coated pit. Mol. Biol. Cell 25, 3595–3609 (2014).

53. Jumper, J. et al. Highly accurate protein structure prediction with AlphaFold. Nature 596, 583–589 (2021).

54. Baek, M. et al. Accurate prediction of protein structures and interactions using a three-track neural network. Science 373, 871–876 (2021).

55. Levental, I. & Lyman, E. Regulation of membrane protein structure and function by their lipid nano-environment. Nat. Rev. Mol. Cell Biol. 24, 107–122 (2023).

56. Virtanen, P. et al. SciPy 1.0: fundamental algorithms for scientific computing in Python. Nat. Methods 17, 261–272 (2020).

57. Ronneberger, O., Fischer, P. & Brox, T. U-Net: Convolutional Networks for Biomedical Image Segmentation. in Medical Image Computing and Computer-Assisted Intervention (MICCAI) (eds. Navab, N., Hornegger, J., Wells, W. M. & Frangi, A. F.) vol. 9351 234–241 (Springer International Publishing, 2015).

58. Akiba, T., Sano, S., Yanase, T., Ohta, T. & Koyama, M. Optuna: A Next-generation Hyperparameter Optimization Framework. in Proceedings of the 25th ACM SIGKDD International Conference on Knowledge Discovery & Data Mining - KDD ‘19 2623–2631 (ACM Press, 2019). doi:10.1145/3292500.3330701.

59. Alex, F., Alex, G., Bertr, RE. Gramfortinria. F., Bertr, T. & Thirion. Scikit-learn: Machine Learning in Python.

60. Kang, Y.-L. et al. Inhibition of PIKfyve kinase prevents infection by Zaire ebolavirus and SARS-CoV-2. Proc Natl Acad Sci USA 117, 20803–20813 (2020).

61. Bohr, F. et al. Enhanced hexamerization of insulin via assembly pathway rerouting revealed by single particle studies. Commun. Biol. 6, 178 (2023).

62. Østergaard, M., Mishra, N. K. & Jensen, K. J. The ABC of insulin: the organic chemistry of a small protein. Chem. Eur. J 26, 8341–8357 (2020).

