## Supplementary figures for "Deep learning assisted single particle tracking for automated correlation between diffusion and function"

**Supporting Information**

**Materials**

For virus experiments in LLSM: Dulbecco modified Eagle's medium (DMEM) supplemented with 4.5 g/liter glucose, L-glutamine, and sodium pyruvate (catalog number 10-013-CV; Corning, Inc.), minimum essential media (MEM) with L-glutamine (catalog number 10-010-CV; Corning, Inc.), Media 199 (catalog number 11150067, Thermo Scientific), OptiMEM media (catalog number 31985070, Life Technologies), FluoroBriteTM DMEM media (catalog number A18967-02, Life Technologies), trypsin/EDTA (catalog number 25-053-CI, Mediatech), fetal bovine serum (FBS) (catalog number S11150H; Atlanta Biologicals), HEPES (catalog number HOL06, Caisson Labs), dimethyl sulfoxide (DMSO) (catalog number 26855; Sigma-Aldrich), Toluene (catalog number 34866, Sigma-Aldrich), dichloromethane (catalog number 270997, Sigma-Aldrich), ethanol (catalog number TX89125172HU, VWR), round cover glass number 1.5 (25 mm, catalog number C8-1.5H-N; Cellvis), polydimethylsiloxane (PDMS) (Sylgard, catalog number DC4019862; Krayden), isopropyl alcohol (catalog number 9080-03/MK303108; VWR), potassium hydroxide (catalog number 484016; Sigma-Aldrich), Aprotinin from bovine lung (catalog number A1153, Sigma-Aldrich), wheat germ agglutinin (WGA)-Alexa Fluor647 (catalog number W32466; Invitrogen), paraformaldehyde (catalog number P6148; Sigma-Aldrich), di-basic sodium phosphate (catalog number BP329-1; Thermo Fisher Scientific), monobasic potassium phosphate (catalog number P5379; Sigma- Aldrich), sodium chloride (catalog number SX0420-5; EMD Millipore), poly-D-lysine (catalog number P6403, Sigma-Aldrich), potassium chloride (catalog number P217-500; Thermo Fisher Scientific), Tris (catalog number T-400-500; Goldbio), EDTA (catalog number E5134; Sigma- Aldrich), sucrose (catalog number S0389;Sigma-Aldrich), fetal calf serum (FCS) (catalog number SH30073.03; HyClone), bovine serum albumin (GE Healthcare), L-glutamine (catalog number G7513; Sigma), Triton X-100 (catalog number 28314; Thermo Fisher). For Insulin experiments on SDCM all reagents are of commercial, analytical grade purchased from Sigma-Aldrich Denmark, unless otherwise stated. ATTO655-NHS (ATTO-TEC). Recombinant human insulin (Thermo Fisher Scientific). MilliQ water was used for aqueous preparations. 10 mM HEPES in HBSS buffer[1].

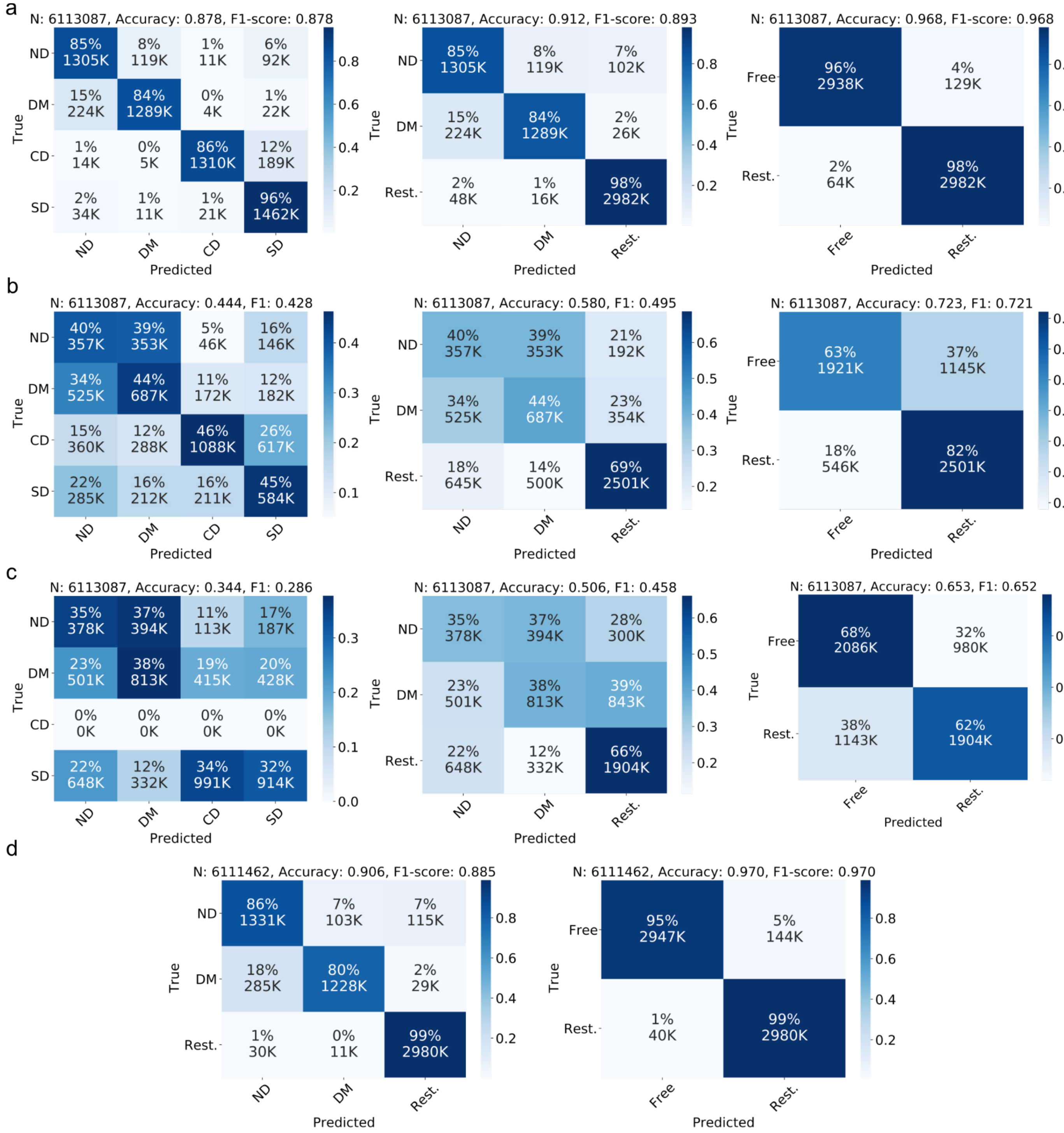


**Fig S1. Comparison of classification accuracy of DeepSPT with LSTM and rolling MSD and effects of combining classes. a,** Confusion matrices of DeepSPT predictions per frame on 2D simulated test set trajectories, Data displayed for either 4 diffusional states, or 3 states normal, directed and confined/subdiffusive, or 2 states normal/directed, confined/subdiffusive. **b,** Confusion matrices of attention BiLSTM[2] predictions per frame on 2D simulated test set trajectories. Data for for 4, 3, 2 diffusional states as above **c,** Confusion matrices of rolling MSD[3,4] predictions per frame on 2D simulated test set trajectories (predicts three classes as it bases predictions on the alpha exponent in a MSD fit based on (subdiffusive alpha)<(normal alpha)<(directed alpha)) Data for for 4, 3, 2 diffusional states as above. **d,** DeepSPT on 3D test set trajectories when combining classes into three and two respectively.

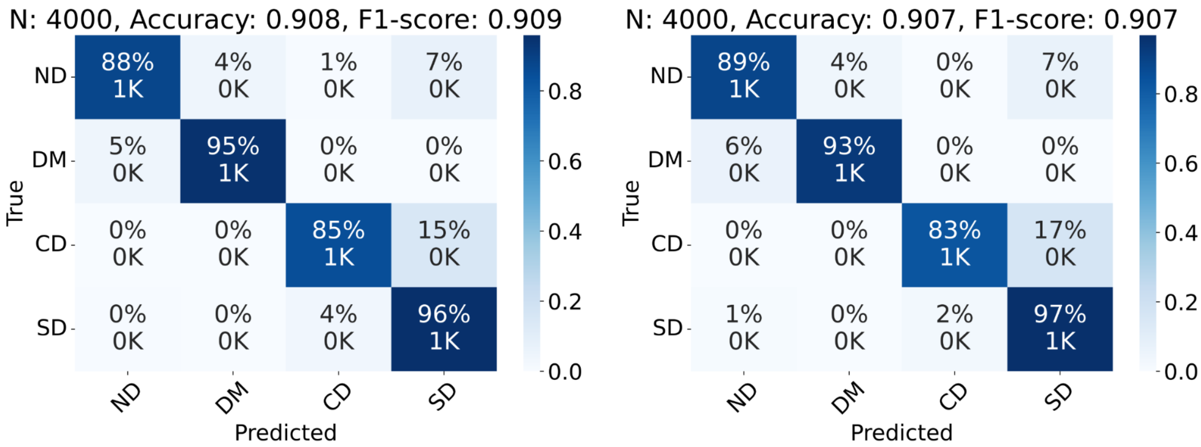


**Fig S2. Confusion matrices for DeepSPT in 2D and 3D for homogeneous motion whole-track predictions. Left,** Confusion matrices of DeepSPT predictions on 2D simulated test set trajectories of homogeneous motion, i.e. predicting the single behavior type exhibited by each trajectory (N=4000). **Right,** Confusion matrices of DeepSPT predictions on 3D simulated test set trajectories of homogeneous motion (N=4000).

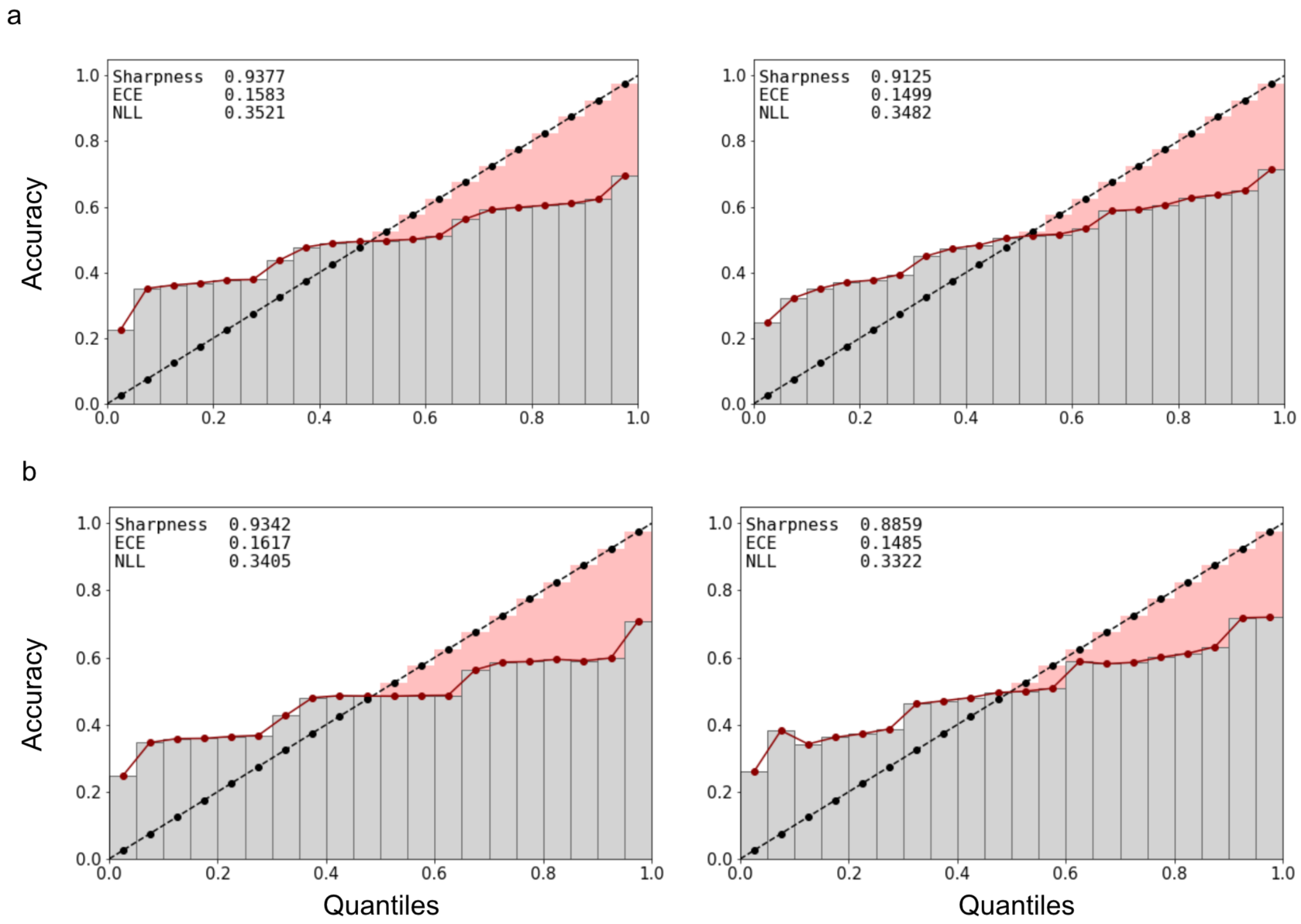


**Fig. S3. Temperature scaling for improving reliability of prediction confidence estimates by DeepSPT towards improved model interpretability and transparency. a,b,** Reliability diagram visualizing model's uncertainty calibration before and after temperature scaling for DeepSPT in 3D and 2D respectively with left figure showing before and right figure showing after uncertainty calibration by temperature scaling[5]. Each reliability plot is accompanied by uncertainty calibration measures; Sharpness (higher is better), expected calibration error (ECE, lower is better) and negative log-likelihood loss (NLL, lower is better). During reliability analysis model predictions are sorted by their class confidence scores on the x-axis plotted against their actual group accuracy, where perfect calibration should be a diagonal line i.e a set predictions which a 10% class probability score should be accurate 10% of the time to be a good probability measure. Inspection of the reliability diagrams report DeepSPT to produce quite reliable probability estimates inherently, but being underconfident for low class probabilities but turns to be overconfident for higher class probabilities, while temperature scaling brings the calibration closer to perfect calibration resulting in improved probability estimates.

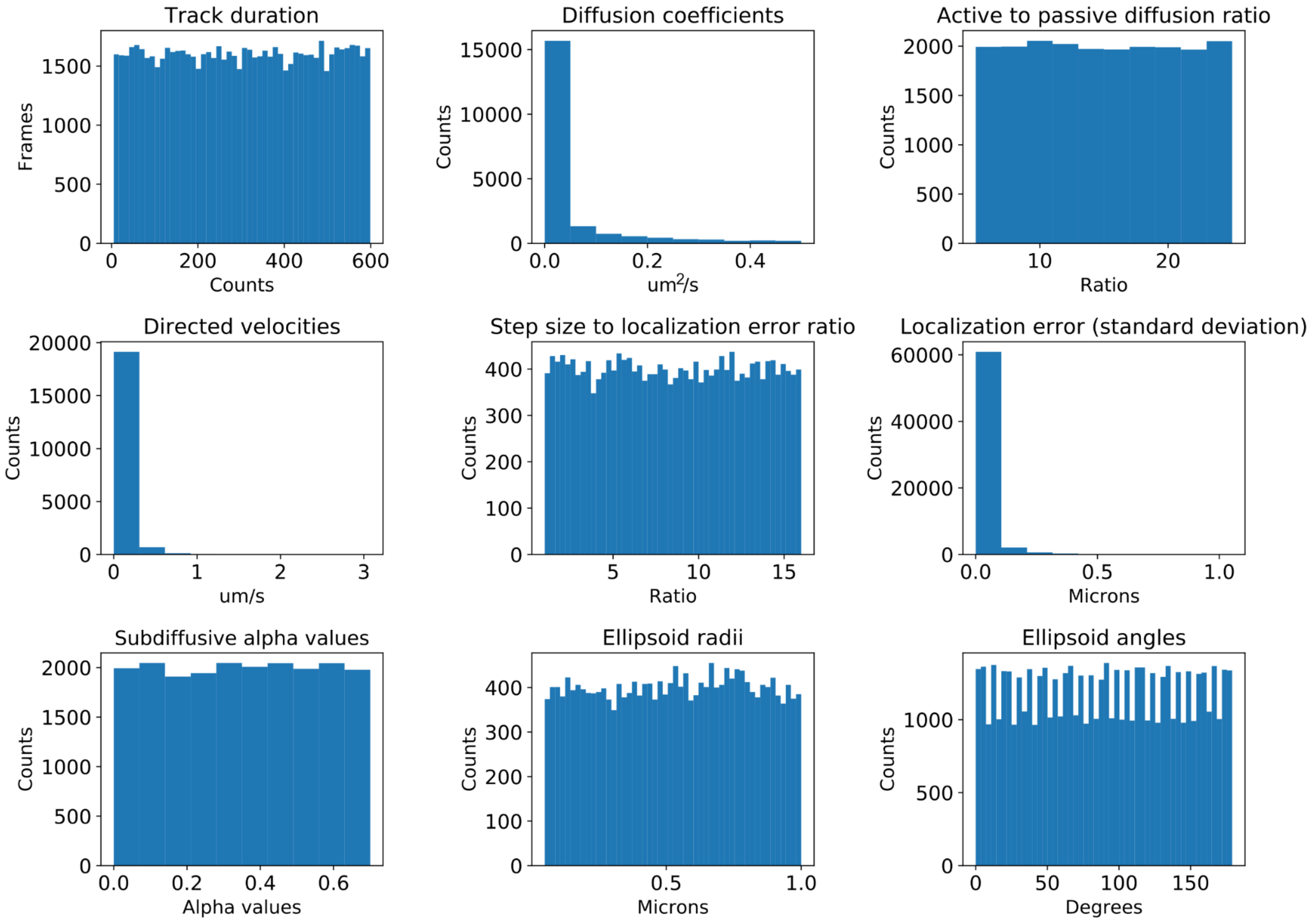


**Fig S4. Distribution of simulation parameters for training and evaluation of DeepSPT.** Histograms showing the distribution of parameters used for stochastic simulation of diffusion. Diffusion coefficients (loguniform), step size to localization error ratio (uniform), localization error (follows diffusion coefficients and localization error ratio) and track durations (uniform) are shared across diffusion types. Active diffusion to passive diffusion ratio (uniform) and directed velocities (follows diffusion coefficients and active-passive ratio) are for simulated directed motion. Subdiffusive alpha values (uniform) are for subdiffusive motion to describe motion persistence while ellipsoid parameters (radii and angles both uniform) are for defining the confinement area of confined motion.

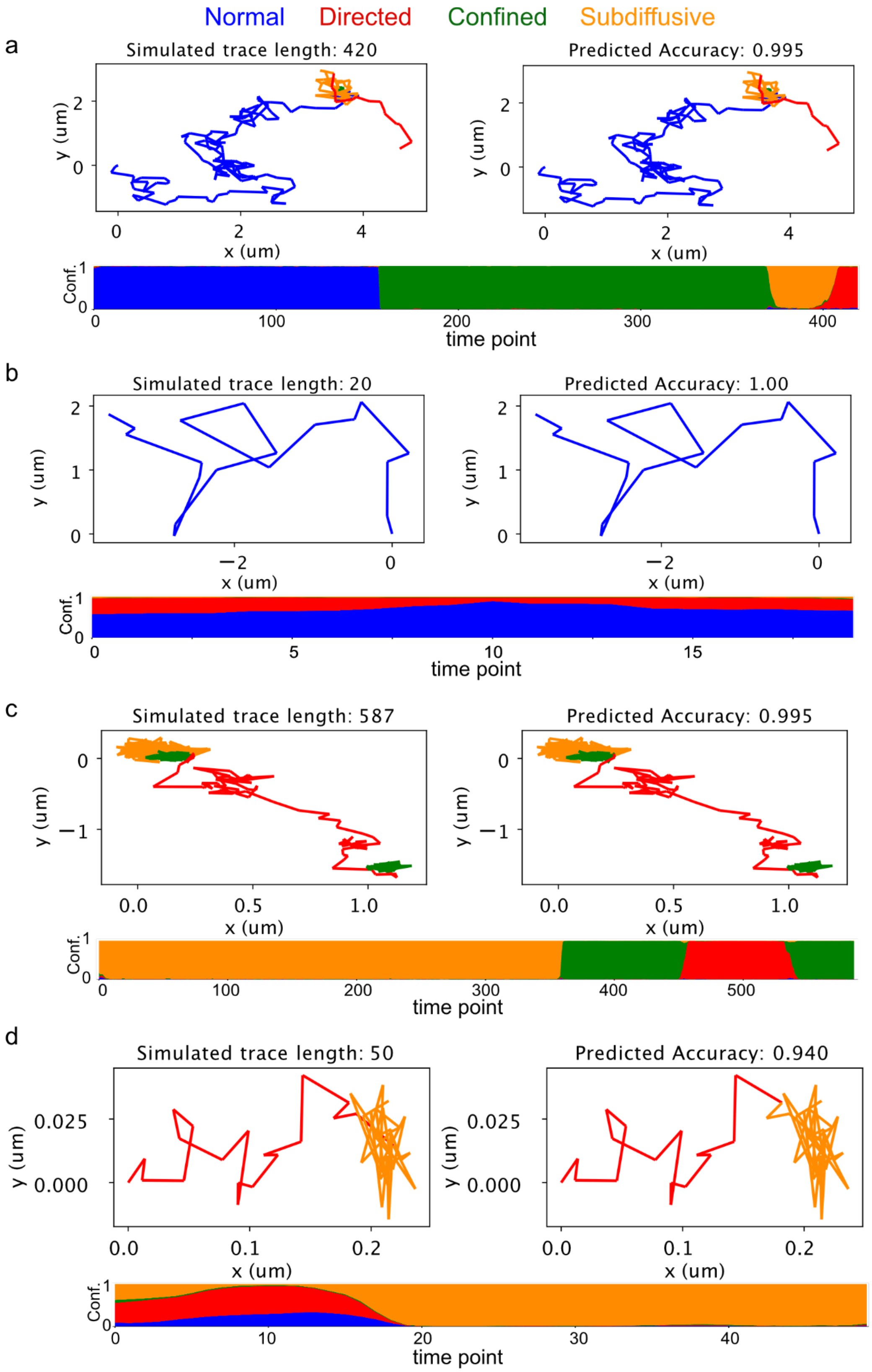


**Fig S5.** Representative examples of Deep-SPT test set predictions on 3D simulated trajectories (**a, b, c, d**). In each panel, raw trace on the left, their prediction on the right and the temporal evolution of temperature scaled behavior type confidence estimates on the bottom.

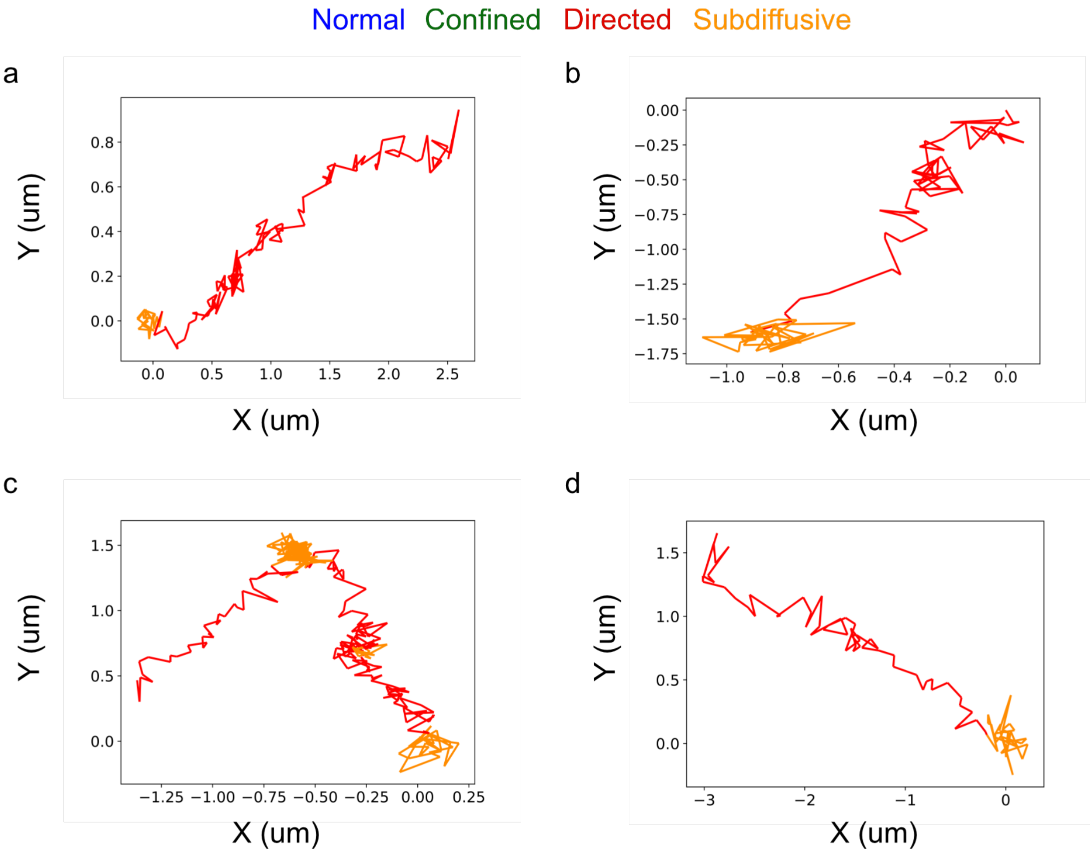


**Fig. 6: Examples of directed behavior exhibited by 2D tracking of human Insulin.** Four panels (**a**,**b**,**c**,**d**) each showing a track identified by DeepSPT to contain directed behavior indicative of active transport processes of human insulin inside HeLa cells.

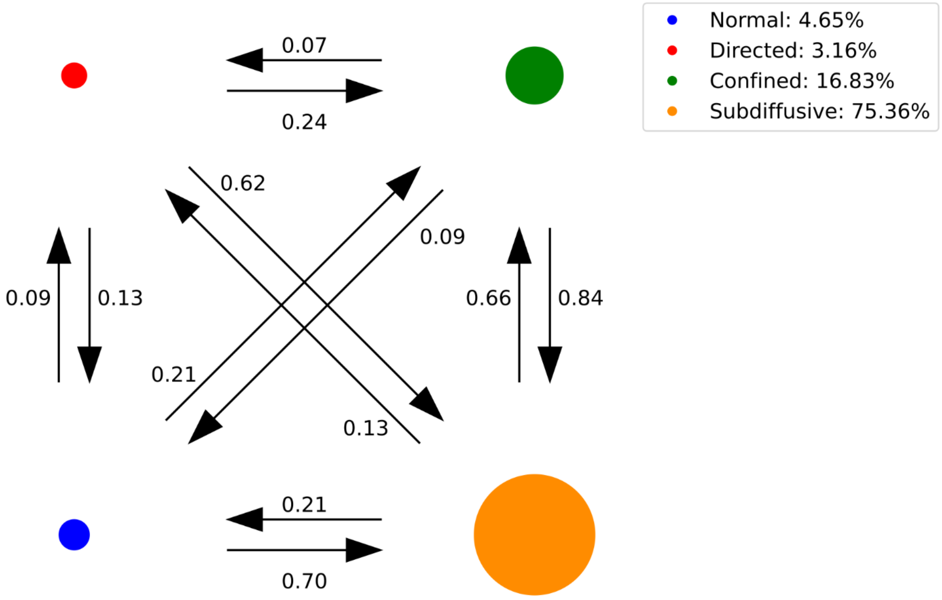


**Fig. S7: Transition likelihoods of human insulin between the 4 identified diffusional states.** The transitions found by DeepSPT between diffusional states normalized so transitions from a given state sum to one to reflect the transition probability for a given diffusional state. Inspection of transition probabilities indicate human insulin movement inside cells favors transitions towards subdiffusive behavior. Legend shows the total occupancy per diffusional state clearly showing subdiffusive behavior being favored by human insulin inside cells but does show directed, characteristic of active transport, and free diffusion does occur. The observed directed motion is indicative of human insulin being actively transported inside cells.

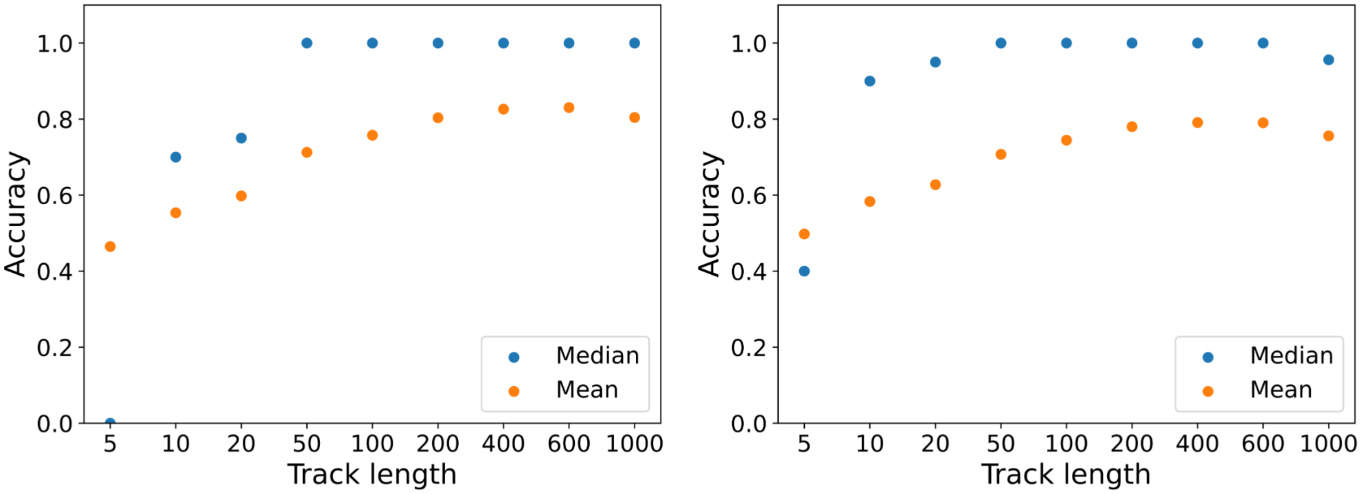


**Fig. S8. Investigating DeepSPT's robustness to various sets of simulation parameters for the homogeneous diffusing test set in 3D.** Investigation of model generalizability and limitations for key simulation parameters for traces containing 1 diffusional types: **a**, 3D Duration of track. **b**, 2D Duration of track. Median accuracy and mean accuracy are track-level metrics providing descriptive statistics for the distribution track-level accuracies for each test set track (N=20000). Flattened accuracy measures the accuracy for all frame-level predictions inside each trajectory pooled together.

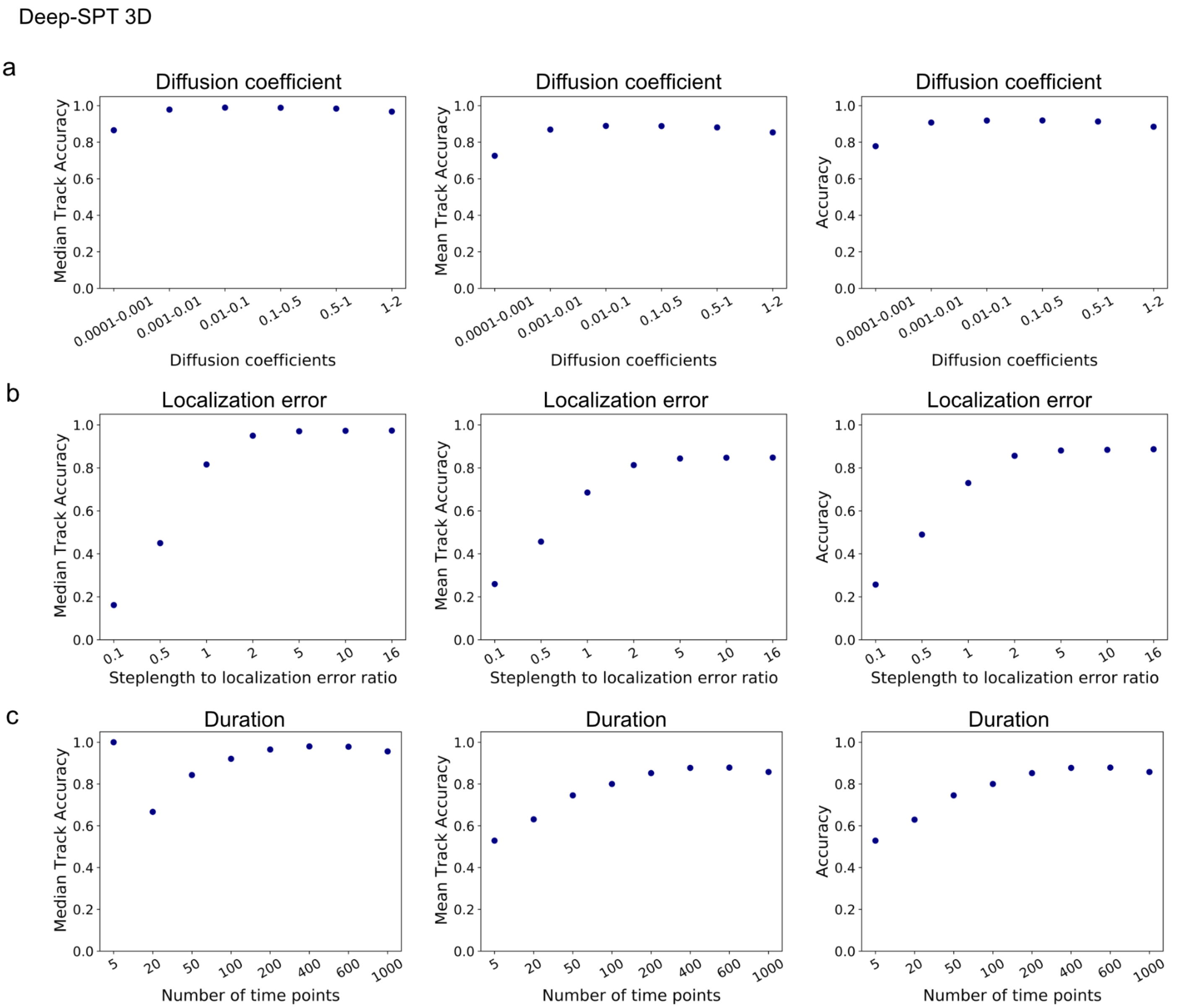


**Fig. S9. Investigating DeepSPT's robustness to various sets of simulation parameters for the heterogeneous diffusing test set in 3D.** Investigation of model generalizability and limitations for key simulation parameters for traces containing all 4 diffusional types at random (see Methods for test set elaboration): **a**, Varying ranges of diffusion coefficients (D) for simulated trajectories (varying D is equivalent to varying temporal resolution for observation of a diffusing particle due to scale invariance of diffusion). **b**, step length to localization error ratio, i.e ratio of contribution to displacements from actual diffusion and localization error respectively where a value of 1 signifies an equal contribution and >1 signifies actual diffusion being larger. **c**, Duration of track. Median accuracy and mean accuracy are track-level metrics providing descriptive statistics for the distribution track-level accuracies for each test set track (N=20000). Flattened accuracy measures the accuracy for all frame-level predictions inside each trajectory pooled together.

Deep-SPT 2D

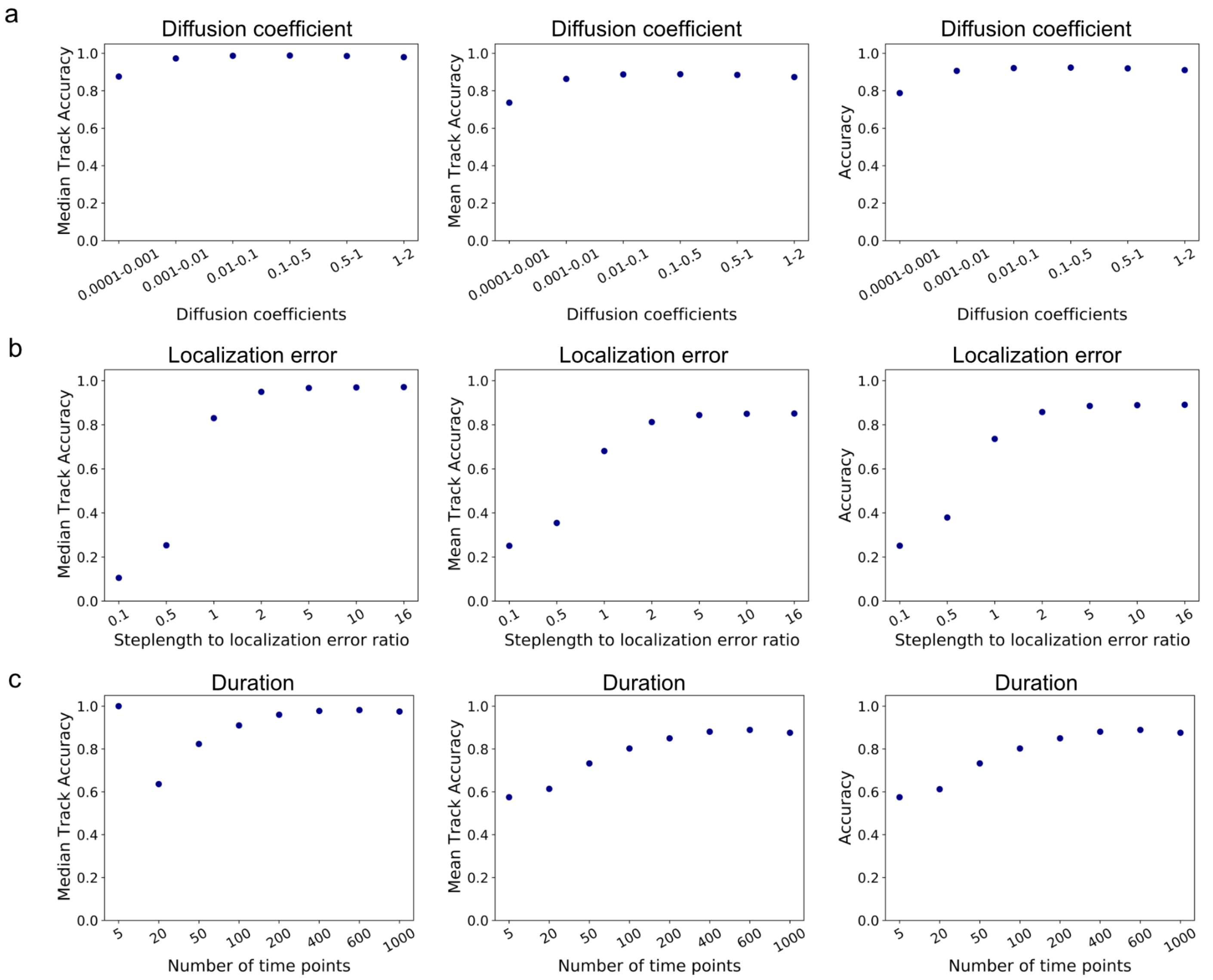


**Fig. S10. Investigating DeepSPT's robustness to various sets of simulation parameters for the heterogeneous diffusing test set in 2D.** Investigation of model generalizability and limitations for key simulation parameters for traces containing all 4 diffusional types at random (see Methods for test set elaboration): **a**, Varying ranges of diffusion coefficients (D) for simulated trajectories (varying D is equivalent to varying temporal resolution for observation of a diffusing particle due to scale invariance of diffusion). **b**, step length to localization error ratio, i.e ratio of contribution to displacements from actual diffusion and localization error respectively where a value of 1 signifies an equal contribution and >1 signifies actual diffusion being larger. **c**, Duration of track. Median accuracy and mean accuracy are track-level metrics providing descriptive statistics for the distribution track-level accuracies for each test set track (N=20000). Flattened accuracy measures the accuracy for all frame-level predictions inside each trajectory pooled together.

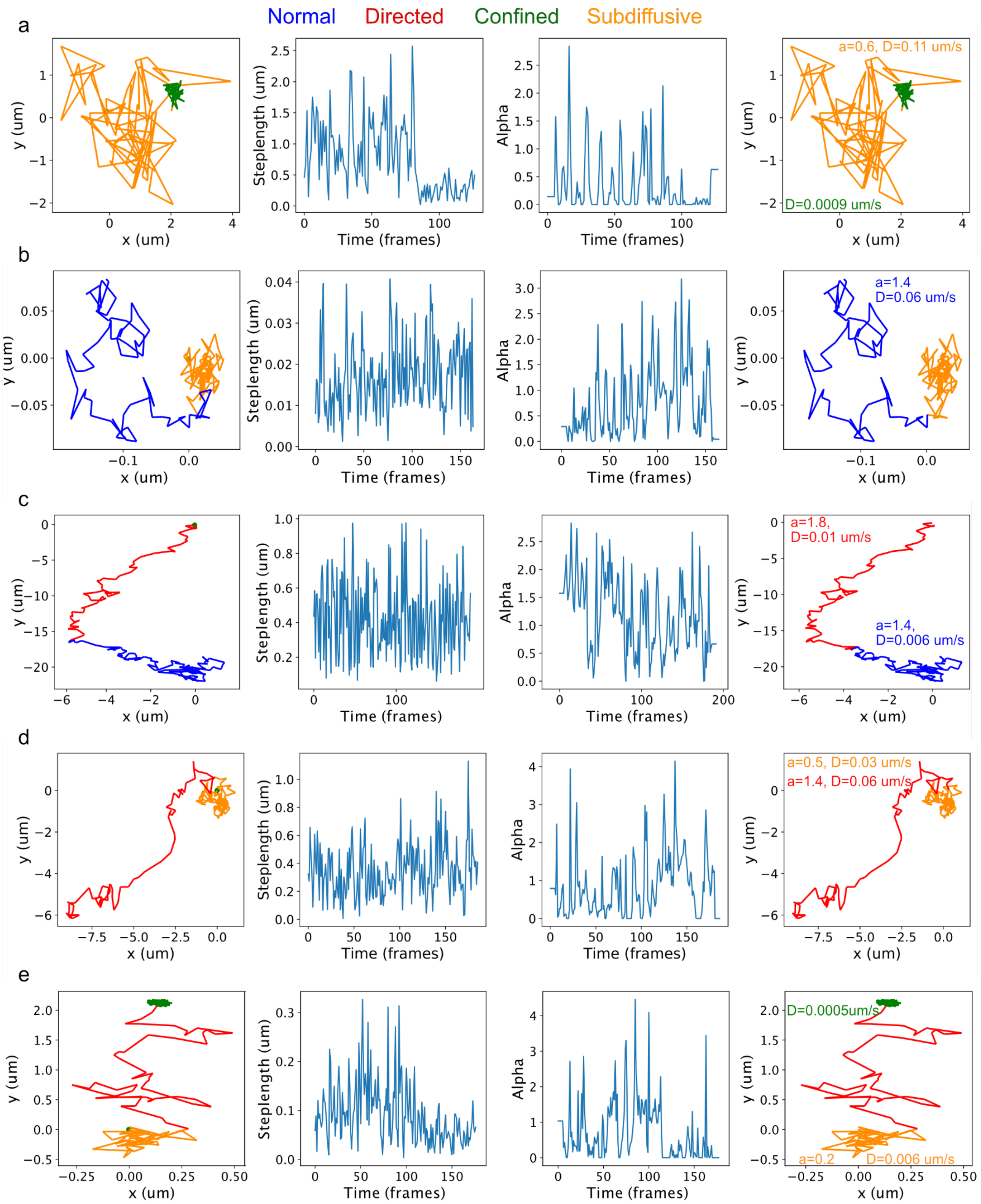


**Fig. S11. Simulated trajectories along conventional temporal diffusional analysis of diffusional metrics and DeepSPT.** Five (**a-e**) rows showing five heterogeneous diffusing trajectories with columns from left to right corresponding to: Ground truth simulation (green dot indicates startpoint), step length time series, their respective MSD analysis, and DeepSPT output. Step length analysis plots the time evolution of step lengths. MSD analysis[3,4] implements a rolling MSD = $2 \cdot dim \cdot D \cdot t^{\alpha}$ producing an alpha estimate for each frame. DeepSPT's outputs temporal diffusional behavior segmentation (normal, directed, confined, subdiffusive) along a set of 40 descriptive features (see Methods). The times series of step lengths and alphas emphasize the challenges faced by HMM and MSD based methods in having to determine discrete underlying states in highly complex and with large variation in the reported diffusional metric.

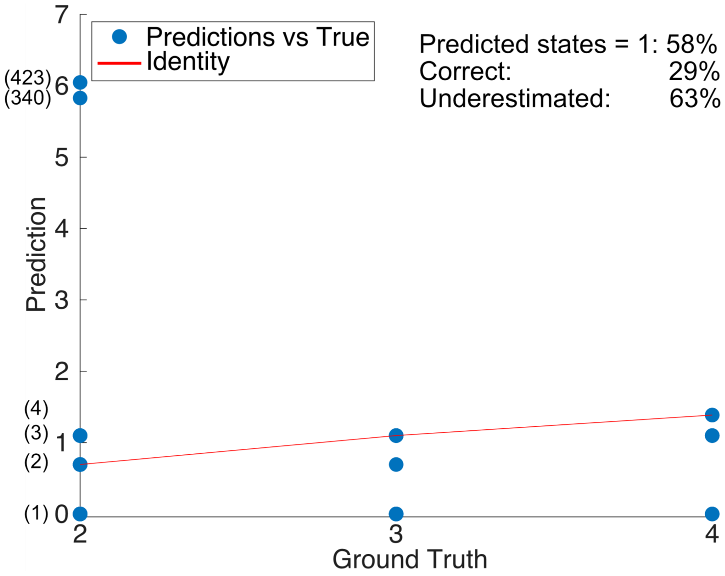


**Fig S12. Segmentation evaluation of HMM-Bayes(ref)**. HMM-Bayes[6] identifies K states of either directed or diffusive within the displacement statistics of individual trajectories the translation of displacement states.HMM-Bayes was asked to search up to K=four different states on randomly selected trajectories with more than 1 diffusional behavior state from the test set utilized in fig. S1. Plot shows the number of states identified by HMM-Bayes versus the actual number of states. The y- axis has been logarithm-transformed for visualization purposes while showing the actual number of states in parenthesis. Overall HMM-Bayes identifies the correct number of states in 29% of the test set even when the maximum number of underlying states was defined a priori by the user. In addition, summary statistics reveal that HMM-Bayes identifies 1 state in 58% of the cases even if no trajectories only contain 1 state and generally underestimates the number of states in 63% of the cases. The required computational time was ~10-60 minutes per trajectory compared to milliseconds by DeepSPT.

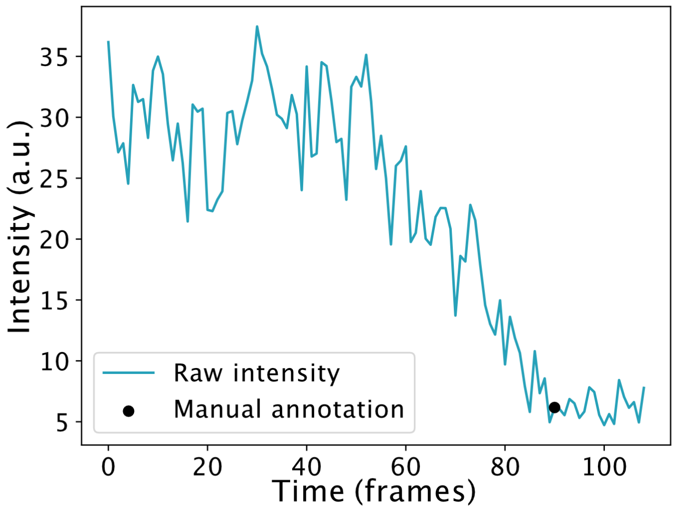


**Fig. S13: a, Example of manual annotation of intensity loss of rotavirus.** Escape into the cytosol of monochromatic rotavirus labelled on free lysines by Atto560 is associated with an intensity drop due to the shedding of the outer layer viral protein VP7[7–9]. Intensity loss is defined manually as the time point at which the intensity drop has seized to fall and the intensity remains constant.

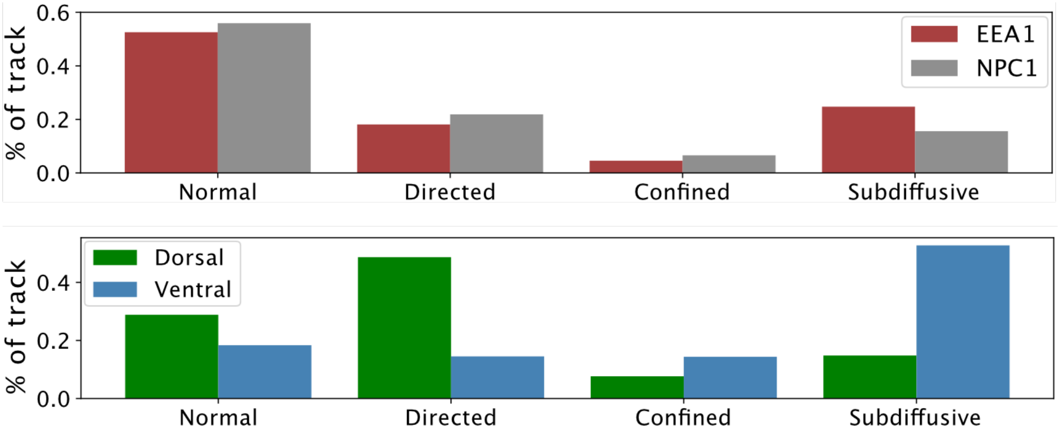


**Fig. S14: Diffusional behavior propensities as predicted by DeepSPT.** Top panel: EEA1- and NPC1- positive endosomes data. Bottom panel: clathrin AP2 adaptor complex data.

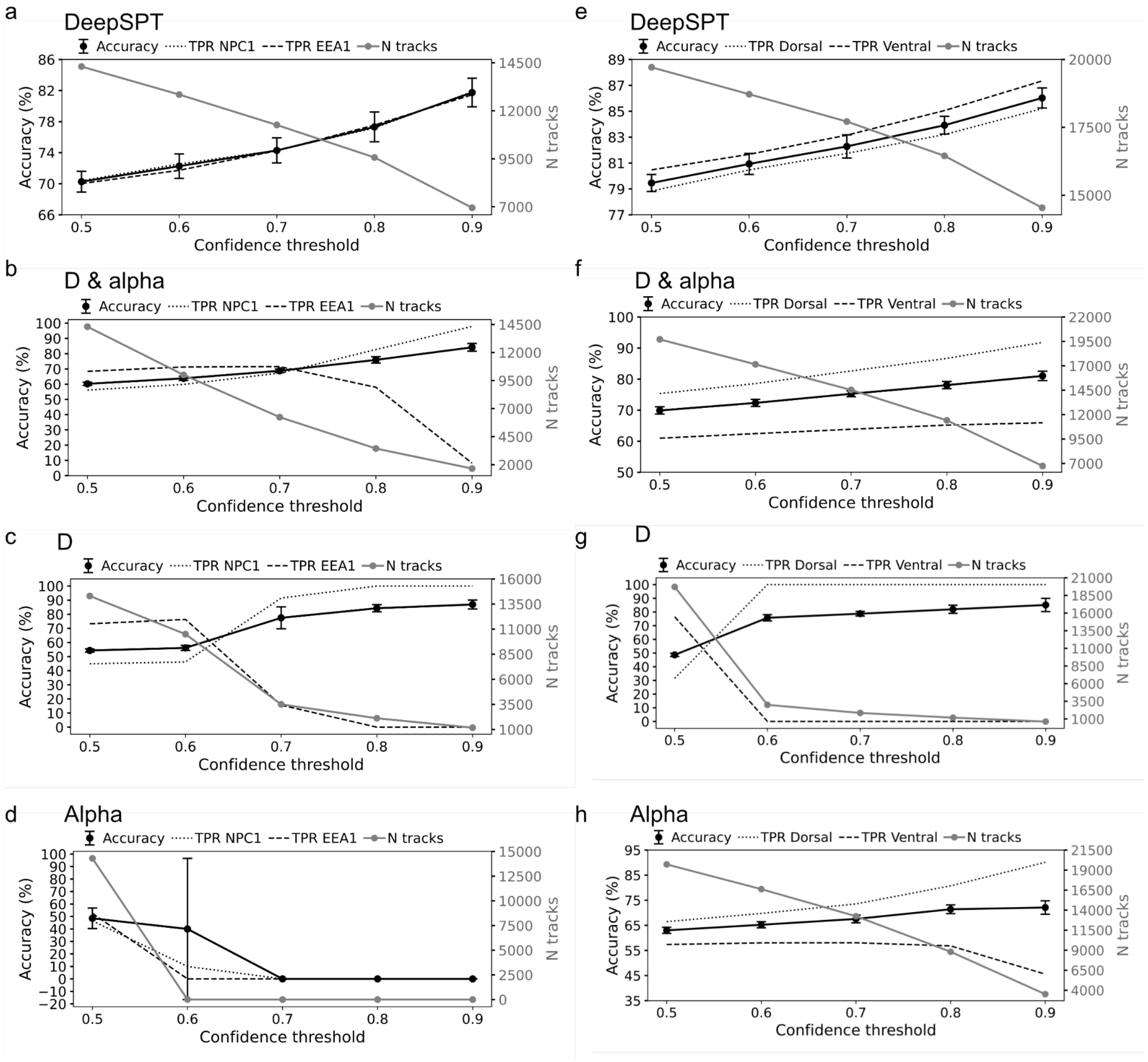


**Fig. S15: Accuracy and number of tracks versus confidence threshold in 10-fold stratified cross-validation.** Akin to main figures 4c and 4h. All panels depict twin axes plot of accuracy (left) and true positive rate (TPR) as well as number of track (right) versus confidence threshold (see Methods). **a-d**: EEA1- and NPC1- positive endosomes data. **e-h**: clathrin AP2 adaptor complex data. Each panel shows the results of either DeepSPT, D & alpha, D, or alpha.

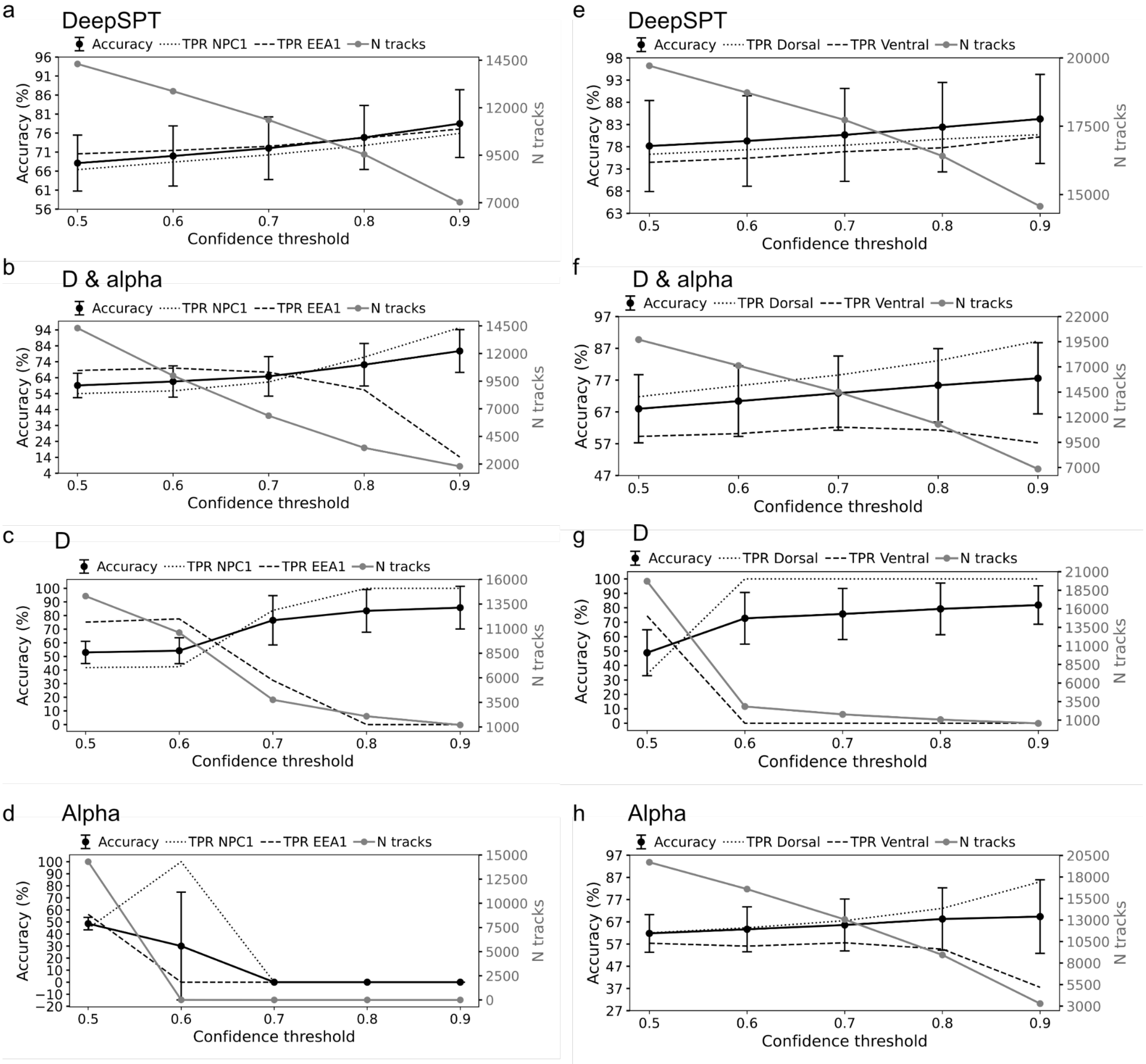


**Fig. S16: Accuracy and number of tracks versus confidence threshold in leave-one-cell-out cross-validation.** Akin to main figures 4c and 4h now leaving an experiment out to show generalization to trajectories in cells not seen by the classifier during training. All panels depict twin axes plot of accuracy (left) and true positive rate (TPR) as well as number of track (right) versus confidence threshold (see Methods). **a-d**: EEA1- and NPC1- positive endosomes data. **e-h**: clathrin AP2 adaptor complex data. Each panel shows the results of either DeepSPT, D & alpha, D, or alpha.

Bibliography

1. Bohr, F. *et al.* Enhanced hexamerization of insulin via assembly pathway rerouting revealed by single particle studies. *Commun. Biol.* **6**, 178 (2023).
2. You, B. & Yang, G. Attention-based LSTM for Motion Switching Detection of Particles in Living Cells. in *2021 International Joint Conference on Neural Networks (IJCNN)* 1–6 (IEEE, 2021). doi:10.1109/IJCNN52387.2021.9533629.
3. Michalet, X. Mean square displacement analysis of single-particle trajectories with localization error: Brownian motion in an isotropic medium. *Phys. Rev. E Stat. Nonlin. Soft Matter Phys.* **82**, 041914 (2010).
4. Arcizet, D., Meier, B., Sackmann, E., Rädler, J. O. & Heinrich, D. Temporal analysis of active and passive transport in living cells. *Phys. Rev. Lett.* **101**, 248103 (2008).
5. Guo, C., Pleiss, G., Sun, Y. & Weinberger, K. Q. On Calibration of Modern Neural Networks. *arXiv* (2017) doi:10.48550/arxiv.1706.04599.
6. Monnier, N. *et al.* Inferring transient particle transport dynamics in live cells. *Nat. Methods* **12**, 838–840 (2015).
7. Salgado, E. N., Garcia Rodriguez, B., Narayanaswamy, N., Krishnan, Y. & Harrison, S. C. Visualization of Calcium Ion Loss from Rotavirus during Cell Entry. *J. Virol.* **92**, (2018).
8. Abdelhakim, A. H. *et al.* Structural correlates of rotavirus cell entry. *PLoS Pathog.* **10**, e1004355 (2014).
9. Aoki, S. T. *et al.* Cross-linking of rotavirus outer capsid protein VP7 by antibodies or disulfides inhibits viral entry. *J. Virol.* **85**, 10509–10517 (2011).
